# Biomni: A General-Purpose Biomedical AI Agent

**DOI:** 10.1101/2025.05.30.656746

**Authors:** Kexin Huang, Serena Zhang, Hanchen Wang, Yuanhao Qu, Yingzhou Lu, Yusuf Roohani, Ryan Li, Lin Qiu, Gavin Li, Junze Zhang, Di Yin, Shruti Marwaha, Jennefer N. Carter, Xin Zhou, Matthew Wheeler, Jonathan A. Bernstein, Mengdi Wang, Peng He, Jingtian Zhou, Michael Snyder, Le Cong, Aviv Regev, Jure Leskovec

## Abstract

Biomedical research underpins progress in our understanding of human health and disease, drug discovery, and clinical care. However, with the growth of complex lab experiments, large datasets, many analytical tools, and expansive literature, biomedical research is increasingly constrained by repetitive and fragmented workflows that slow discovery and limit innovation, underscoring the need for a fundamentally new way to scale scientific expertise. Here, we introduce Biomni, a general-purpose biomedical AI agent designed to autonomously execute a wide spectrum of research tasks across diverse biomedical subfields. To systematically map the biomedical action space, Biomni first employs an action discovery agent to create the first unified agentic environment – mining essential tools, databases, and protocols from tens of thousands of publications across 25 biomedical domains. Built on this foundation, Biomni features a generalist agentic architecture that integrates large language model (LLM) reasoning with retrieval-augmented planning and code-based execution, enabling it to dynamically compose and carry out complex biomedical workflows – entirely without relying on predefined templates or rigid task flows. Systematic benchmarking demonstrates that Biomni achieves strong generalization across heterogeneous biomedical tasks – including causal gene prioritization, drug repurposing, rare disease diagnosis, microbiome analysis, and molecular cloning – without any task-specific prompt tuning. Real-world case studies further showcase Biomni’s ability to interpret complex, multi-modal biomedical datasets and autonomously generate experimentally testable protocols. Biomni envisions a future where virtual AI biologists operate alongside and augment human scientists to dramatically enhance research productivity, clinical insight, and healthcare. Biomni is ready to use at https://biomni.stanford.edu, and we invite scientists to explore its capabilities, stress-test its limits, and co-create the next era of biomedical discoveries.

## 1 Introduction

Biomedical research is a key pillar of modern science and medicine, driving discoveries in disease mechanisms, diagnostics, and therapeutics^1–4^. Yet, with the growth in large-scale experiments, data, tools, and literature, progress is increasingly slowed by fragmented, complex workflows that require specialized tools, exhaustive literature reviews, intricate experimental design, and careful statistical modeling^5, 6^. A vast volume of valuable biomedical data sits underutilized^7^, many sophisticated analyses are not conducted, and many connections for past knowledge and literature are not made, not for lack of significance, but because the demand for expert researchers far exceeds the supply. This mismatch between data abundance and limited human bandwidth highlights an urgent need for a fundamentally new approach – one that can effectively scale expertise, streamline workflows, and unlock the full potential of biomedical research.

Recent advances in Artificial Intelligence (AI) have created a paradigm shift, opening the possibility for fundamentally reshaping biomedical research^8^. AI agents have dramatically reshaped fields such as software engineering^9^, law^10^, material science^11^ and healthcare^12^ by automating repetitive tasks, enhancing productivity, and enabling breakthroughs that were previously unimaginable. Given these developments, the question emerges: *Can we build a virtual AI biomedical scientist?* Such a virtual scientist would autonomously tackle diverse biomedical research tasks spanning multiple subfields, unlocking extensive capabilities and fostering novel insights through interdisciplinary integration – an achievement that can radically augment human biologists limited by specialized expertise. Capable of efficiently managing thousands of concurrent tasks, this virtual scientist could dramatically enhance human productivity and accelerate the pace of biomedical discovery.

Previous approaches have largely relied on specialist agentic workflows tailored to narrow biomedical tasks^13–19^, which restricts their capacity to move fluidly and generalize across the full spectrum of biomedical domains, as needed to answer key research questions. Enabling an AI agent to handle a broad range of biomedical tasks introduces substantial technical challenges – most notably, the need to tightly couple advanced reasoning^20^ with the ability to execute highly specialized biomedical actions^21^. Although LLM-based reasoning has seen significant advancements^22^, such LLMs need access to an environment that explicitly defines the biomedical action space, which is inherently diverse, domain-specific, and complex. Moreover, a truly capable system requires an agentic architecture that can natively interact with this biomedical environment – autonomously selecting and composing actions, using its reasoning capabilities to plan and execute diverse tasks without relying on rigid, pre-defined workflows.

Here we present Biomni, a generalist biomedical AI agent purpose-built to automate and advance biomedical research across a wide range of subfields. Acting as a virtual AI biologist, Biomni autonomously formulates novel, testable hypotheses, performs complex bioinformatics analyses, and designs rigorous experimental protocols. To enable this capability, we first constructed a unified and comprehensive biomedical action space by systematically analyzing tens of thousands of biomedical research papers spanning 25 distinct subfields, curated from major bio-literature repositories. From this foundation, we developed an LLM-powered action discovery agent capable of reading papers and extracting key tasks, tools, and databases essential to driving biomedical discoveries. These elements are then selected and implemented into Biomni-E1, the foundational environment that defines the biomedical action space for agentic interaction. Biomni-E1 includes 150 specialized biomedical tools, 105 software packages, and 59 databases. We then designed Biomni-A1, a general-purpose agent architecture capable of flexibly executing a broad spectrum of biomedical tasks by using tools and datasets provided by Biomni E1. Given a user query, the agent first uses a retrieval system to identify the most relevant tools, databases, and software needed. It then applies LLM-based reasoning and domain expertise to generate a detailed, step-by-step plan. Each step is expressed through executable code, enabling precise and flexible compositions of biomedical actions – an essential feature given the domain’s reliance on highly specialized tools and data resources. Unlike traditional function-calling methods, this approach supports the dynamic and complex nature of biomedical workflows. This integrated system allows Biomni not only to solve challenging, large-scale biomedical problems with efficiency, but also to generalize to novel tasks across previously unseen areas of biomedical research.

Rigorous benchmarking demonstrates Biomni’s outstanding performance across established biomedical Q&A benchmarks, and robust generalization performance in eight challenging, realistic scenarios never encountered during development. Additionally, we highlight Biomni’s practical capabilities through three impactful case studies: (1) analyzing 458 files of wearable sensor data to generate novel insights; (2) rapidly performing comprehensive bioinformatics analyses on massive raw datasets, such as single-cell RNA-seq and ATAC-seq data, to generate novel insights and hypotheses; (3) autonomously designing laboratory protocols to assist wet-lab researchers. With Biomni, we introduce the first generation of a scalable, general-purpose biomedical AI agent, setting the stage for an era where virtual AI biologists work alongside human researchers to dramatically accelerate biomedical discovery from basic research to translation.

## 2 Results

### Overview of Biomni

Biomni is a general-purpose biomedical AI agent comprising two main components: Biomni-E1, a foundational biomedical environment with a unified action space, and Biomni-A1, an intelligent agent designed to utilize this environment effectively.

Curating a unified biomedical action space is challenging due to its inherent complexity and vastness. We systematically address this by employing an AI-driven approach (Figure 1a). Specifically, we leveraged the 25 subject categories defined by bioRxiv, selecting the 100 most recent publications per category. An action discovery LLM agent processed each paper sequentially, extracting essential tasks, tools, databases, and software necessary to replicate or generate the described research. This comprehensive set of resources constitutes the essential actions required to perform a large set of biological research tasks.

**Figure 1:**
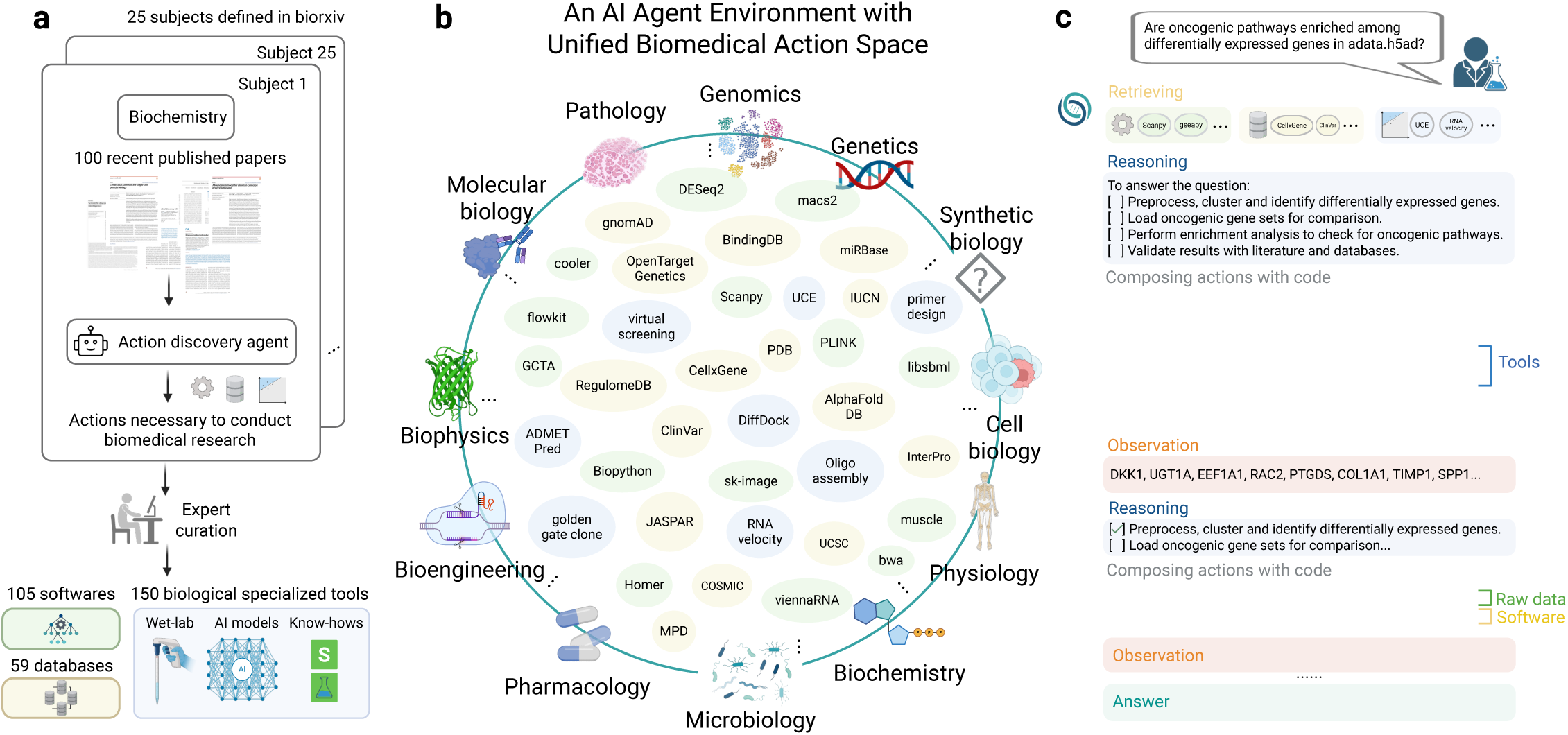
Overview of the unified biomedical action space and agent environment in Biomni. (a) Workflow for systematically curating the unified biomedical action space. Actions necessary to conduct biomedical research were extracted from 2,500 recent bioRxiv publications across 25 biomedical subfields using an AI-driven discovery agent. Extracted actions were rigorously validated and curated by human experts, resulting in the integration of 105 biomedical software tools, 150 specialized biological tools (including wet-lab protocols, AI-driven predictive models, and domain-specific know-how), and 59 comprehensive biomedical databases. (b) Illustration of the unified biomedical action space spanning diverse biomedical subfields such as genetics, genomics, synthetic biology, cell biology, physiology, microbiology, pharmacology, bioengineering, biophysics, molecular biology, and pathology. Representative tools and databases integrated into Biomni’s environment are shown, highlighting its general-purpose capabilities. (c) Example workflow demonstrating Biomni’s reasoning and action composition process to autonomously answer a complex biological question. Biomni retrieves relevant tools based on the user’s query, formulates a structured reasoning plan, and composes executable code to perform comprehensive bioinformatics analyses, iteratively refining its reasoning based on observations until converging on a final, precise answer.

We then curated Biomni-E1, an environment for a biomedical AI agent to perform a wide range of actions (Figure 1b). Identified tools were rigorously verified by human experts, along with corresponding test cases. These tools (Supplementary Table 1-Table 18) were specifically chosen for their non-trivial nature, encompassing complex code, domain-specific know-how, or specialized AI models. Recognizing the inherent flexibility required by biological software, which cannot always be simplified into static functions, we constructed an execution environment preinstalled with 105 widely-used biological software packages (Supplementary Table 23-30), supporting Python, R, and Bash scripts. For database integration, we categorized resources into two distinct groups. The first group consists of massive relational databases accessible via web APIs (e.g., PDB, OpenTarget, ClinVar) (Supplementary Table 19-20). Rather than creating numerous individual retrieval tools, we implemented a unified function per database. Each function accepts natural language queries and internally employs an LLM to parse database schemas and generate executable queries dynamically. Databases without web interfaces were downloaded into a data lake and preprocessed locally into structured pandas DataFrames for seamless integration with the agent, for a total of 59 databases in Biomni-E1 (Supplementary Table 21-22). In summary, Biomni-E1 is the first environment for biomedical AI agent and includes 150 specialized biomedical tools, 105 software, and 59 databases.

To build a general-purpose agent capable of tackling diverse biomedical tasks, we require a specialized agentic architecture – one that avoids hardcoding workflows for each individual task. This led to the development of Biomni-A1, which incorporates several core innovations critical for operating across the biomedical research landscape. First, we introduce an LLM-based tool selection mechanism designed to navigate the complexity and specialization of biomedical tools, dynamically retrieving a tailored subset of resources based on the user’s goal. Second, recognizing that biomedical tasks often require rich procedural logic, Biomni-A1 uses code as a universal action interface – allowing it to compose and execute complex workflows involving loops, parallelization, and conditional logic. Crucially, this approach also enables the agent to interleave calls to software, tools, databases, and raw data operations that do not conform to predefined function signatures-supporting flexible and dynamic integration of heterogeneous resources. Third, the agent adopts an adaptive planning strategy: it formulates an initial plan grounded in biomedical knowledge and iteratively refines it throughout execution, enabling responsive, context-aware behavior. Together, these innovations enable Biomni-A1 to generalize to previously unseen tasks and domains, dynamically composing intelligent actions and interfacing with software, data, and tools in a way that embodies generalist biomedical intelligence (Figure 1c).

### Biomni excels on general biomedical knowledge and reasoning benchmarks

We evaluated Biomni on three challenging multiple-choice benchmarks of general biomedical knowledge and reasoning: Humanity’s Last Exam (HLE)^23^ and LAB-Bench^24^, which includes two key subtasks – DbQA (Database Question Answering) and SeqQA (Sequence Question Answering) (Figure 2a). These tasks span tool use, symbolic reasoning, and structured biological information retrieval – core competencies for any robust biomedical AI agent. To isolate the impact of tool access and agent design, we compared Biomni against six strong general-purpose baselines (details in Supplementary Notes A). Specialized methods^25, 26^ designed for each task is not considered as we aim to compare on generalist performance.

**Figure 2:**
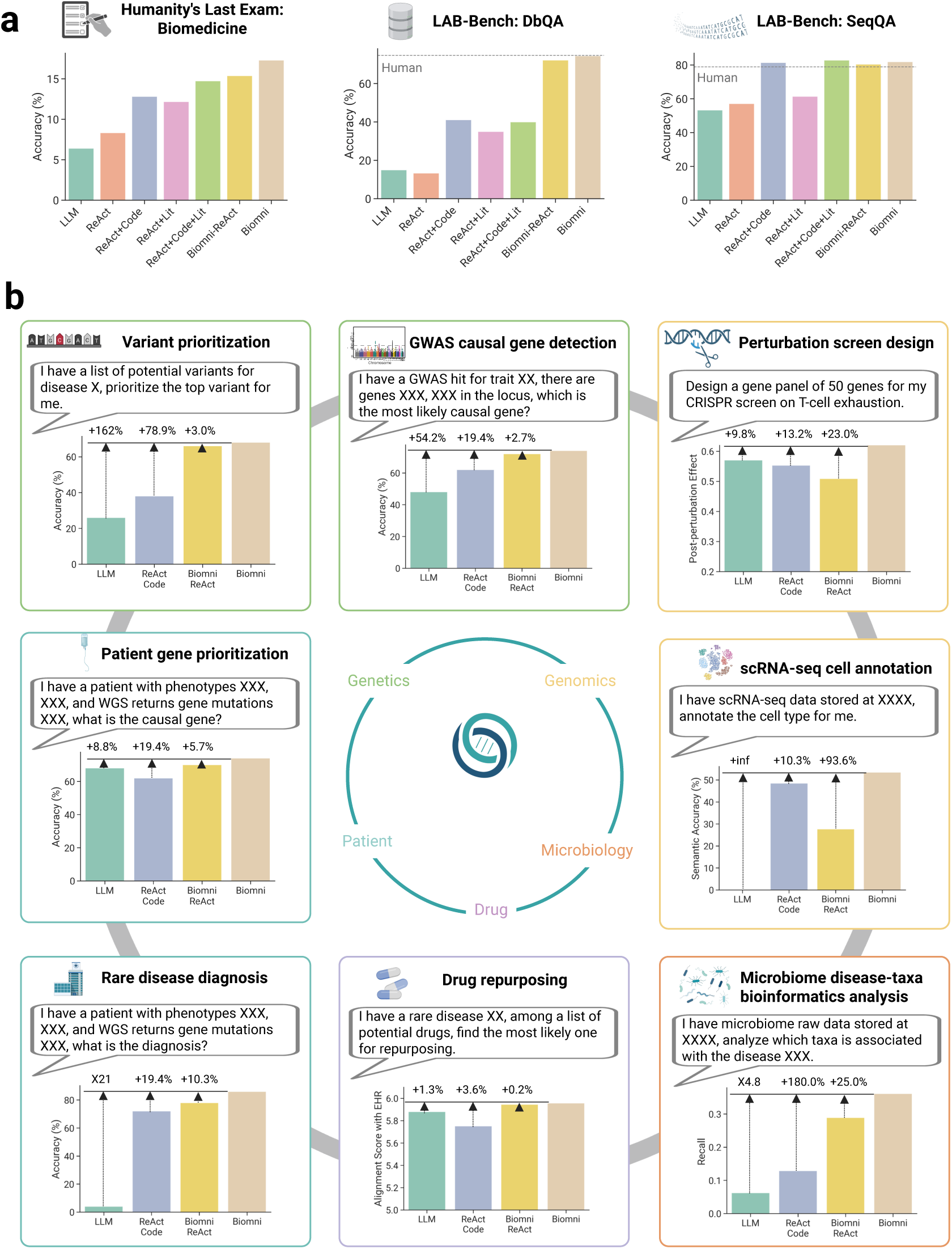
Zero-shot generalization of Biomni across diverse realistic biomedical tasks. (a) Biomni is superior to 6 baselines in Q&A multiple choice benchmarks that broadly evaluate the model’s capability across biomedical fields. (b) Biomni demonstrates robust zero-shot performance across eight previously unseen, real-world biomedical scenarios spanning multiple biomedical subfields, without any task-specific fine-tuning or prompt engineering. Evaluated tasks include variant prioritization and GWAS causal gene detection (genetics and genomics), perturbation screen design (functional genomics, immunology), patient gene prioritization, rare disease diagnosis (clinical genomics), drug repurposing (pharmacology), microbiome disease-taxa bioinformatics analysis (microbiology), and single-cell RNA-seq cell annotation (single-cell biology). Across these diverse scenarios, Biomni consistently outperformed baseline models (Base LLM, ReAct+Code) and specialized environments (Biomni ReAct), highlighting its general-purpose biomedical capabilities and ability to autonomously adapt to new and complex biomedical tasks.

For LAB-Bench, a 45-question development set was used to refine tool and database interfaces, while the final evaluation was conducted on 315 held-out test questions, with performance averaged across three independent runs. We only curated a representative 12.5% subset of the full benchmark due to API cost constraints. In DbQA, which requires structured querying over biological databases, Biomni achieved 74.4% accuracy – matching expert human performance (74.7%) and outperformed all baselines, including the coding agent (ReAct+Code, 40.8%). In SeqQA, which involves reasoning over DNA and protein sequences, Biomni achieved 81.9% accuracy, again exceeding human-level performance (78.8%).

To test true generalization of biomedical knowledge and reasoning *without any development set*, we also evaluated Biomni on a 52-question subset of HLE spanning 14 biomedical subfields – from molecular biology to physiology. Biomni achieved 17.3% accuracy, significantly outperforming the base LLM (6.0%), coding agent (12.8%), and literature agent (12.2%). These results demonstrate Biomni’s ability to generalize across unfamiliar, open-ended biomedical domains without any task-specific adaptation. Additional ablation results are shown in Supplementary Figures 1-2. Performances across each subfield are reported in the Supplementary Figure 3.

### Biomni generalizes to new, real-world biomedical tasks across diverse subfields

To evaluate generalization in realistic research tasks, we curated eight new biomedical benchmarks spanning genetics, genomics, microbiology, pharmacology, and clinical medicine (Figure 2b). Each task was framed to reflect a common, well-defined, but complex real-world biomedical research goal, including: (1) Variant prioritization: Identify the most likely causal variant from a list of potential variants for a trait, requiring reasoning about regulatory functions in non-coding regions. (2) GWAS causal gene detection: Select the most likely causal gene within a locus, demanding fine-grained locus-level inference. (3) CRISPR perturbation screen design: Construct gene panels to maximize post-perturbation effect across a large (⇠20,000 genes) search space. (4) Rare disease diagnosis: Map patient phenotypes and genetic findings to rare disease diagnosis. (5) Drug repurposing: Given a rare disease and a list of candidate drugs, select the best therapeutic match. (6) Single-cell RNA-seq annotation: Assign accurate cell-type labels to individual cell profiles across tissues, species, and platforms. (7) Microbiome disease-taxa analysis: Perform statistical association tests on microbiome datasets to uncover disease-relevant taxa. (8) Patient gene prioritization: Given an individual patient’s genetic profile and phenotype description, identify the most plausible causal gene. We benchmarked Biomni without prompt engineering or task-specific fine-tuning against three baselines: (1) a base LLM (Claude Sonnet 3.7) without tool use, (2) a coding agent with direct function calls and code execution (ReAct+Code), and (3) Biomni-ReAct, an ablation of Biomni that replaces code-based planning with ReAct-style chaining. The complete benchmark constructions are described in Methods, with detailed performance comparisons in Supplementary Notes B.

Across all tasks, Biomni outperformed the base LLM by an average relative performance gain of 402.3%, the coding agent by 43.0%, and its own ablated variant Biomni-ReAct by 20.4%. These findings highlight the importance of code-centric planning and environment grounding, enabling Biomni to compose precise, flexible, and context-aware actions. For each benchmark, we further analyzed the execution trajectories, identifying commonly invoked tools, software, and datasets, as detailed in Supplementary Figures 6-16. These trajectories provide insight into the complexity and structure of agent behavior across tasks. On average, Biomni executes between 6 and 24 distinct steps per task, involving combinations of 0-4 specialized tools, 1-8 software packages, and 0-3 unique data lake items. The agent interleaves data extraction, search/retrieval, reasoning, and computational analyses (Supplementary Figure 8) – reflecting a workflow pattern that mirrors how human scientists alternate between retrieving knowledge and generating new insights. Resource usage varies by task type: information synthesis tasks, such as CRISPR perturbation screen design and GWAS causal gene identification, rely heavily on database queries (e.g., KEGG, Reactome) and literature search (e.g., PubMed, Google), whereas bioinformatics analysis tasks like microbiome profiling and single-cell annotation involve minimal database use but extensive code execution with software libraries such as scanpy.

### Biomni jointly analyzes 458 wearable sensor files to generate physiological hypotheses

To evaluate Biomni’s performance in real-world biomedical workflows, we invited scientists to apply it directly to their own research questions. In this case study, a researcher used Biomni to analyze 458 Excel files containing months-long wearable sensor data (continuous glucose monitoring (CGM) and body temperature) from 30 participants. The data were highly heterogeneous: file formats varied, annotations were inconsistent, and participants exhibited substantial variability (Figure 3a). The researcher posed an open-ended question: Can we uncover biologically meaningful thermogenic patterns?

**Figure 3:**
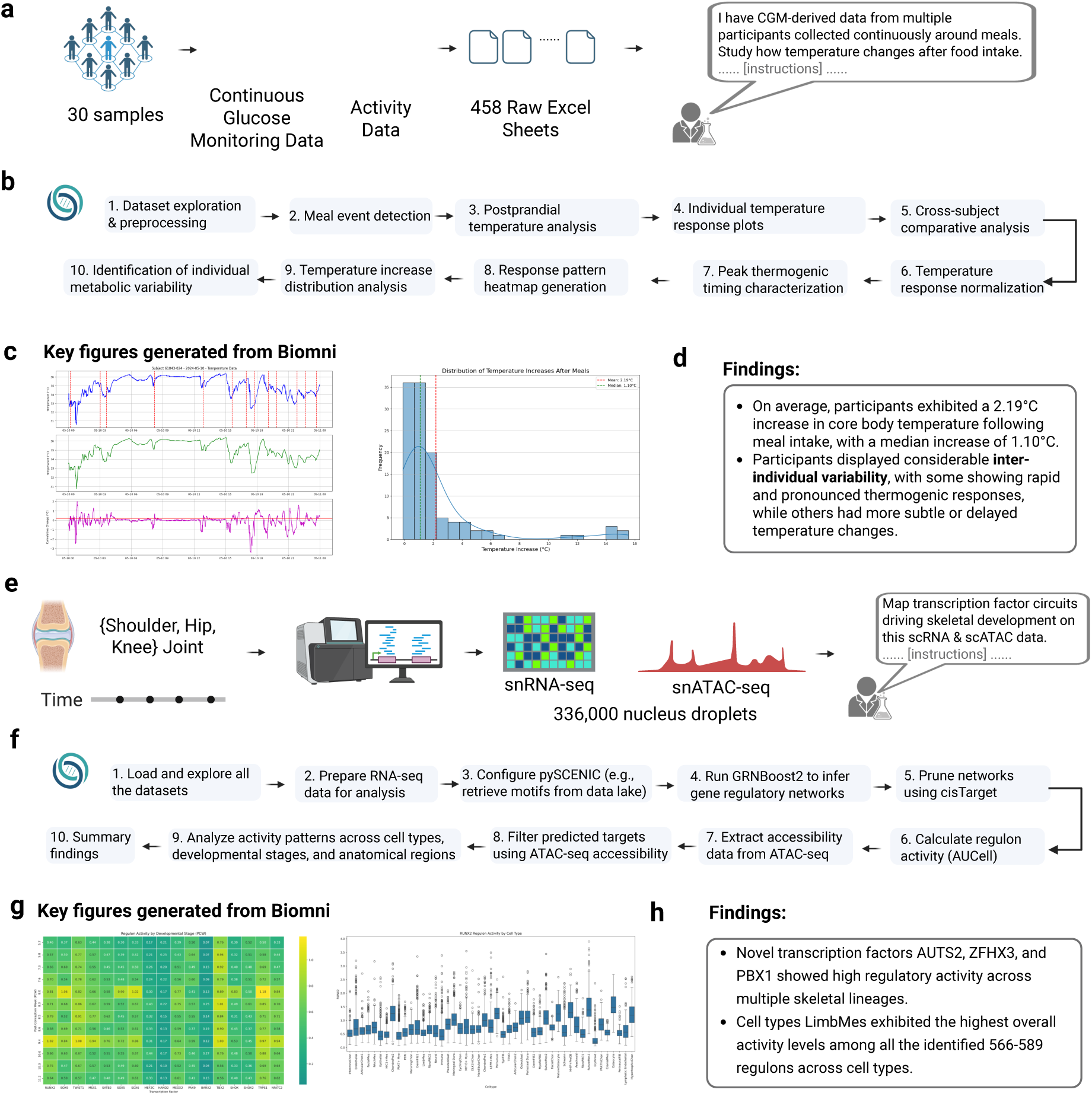
Biomni autonomously executes complex multi-modal biomedical analyses to generate hypothesis. (a-d) Biomni rapidly analyzed CGM-derived thermogenic responses data and activity data from 30 individuals, comprising 458 raw Excel sheets. (b) Workflow demonstrating Biomni’s autonomous execution of data preprocessing, meal event detection, postprandial temperature analysis, and thermogenic response characterization. (c) Representative individual temperature-response plots and temperature increase distribution following meals, automatically generated by Biomni. (d) Summary of unique biological findings identified by Biomni, including significant increases in core body temperature post-meal intake (average 2.19C, median 1.10C), and notable inter-individual variability in thermogenic responses. (e-h) Biomni autonomously analyzed single-cell multiomics data from approximately 336,000 nucleus droplets, combining single-nucleus RNA (snRNA-seq) and single-nucleus ATAC sequencing (snATAC-seq) across human embryonic joint development (shoulder, hip, knee). (f) A detailed workflow diagram showing Biomni’s 10-step analysis pipeline for gene regulatory networks with multiomics. (g) Two key figures generated from Biomni: Left panel shows a heatmap of regulator activity by developmental stage, with color intensity indicating activity levels. Right panel displays a boxplot of RUNX2 regulon activity by cell type, showing variation in expression across different cell populations. (h) Key findings from the GRN analysis: 1) Novel transcription factors (AUTS2, ZFHX3, and PBX1) showing high regulatory activity across multiple skeletal lineages despite no previous association with skeletal development, and 2) Across the 566-589 regulons recovered, limb mesenchyme cells display the highest mean regulonactivity score, underscoring their prominent role in skeletal transcriptional control.

Biomni autonomously generated and executed a 10-step analysis pipeline (Figure 3b), inferring meal events from glucose spikes, extracting pre/post meal temperature windows, normalizing across individuals, and synthesizing population-level trends. Crucially, after completing the pipeline, the agent delivered a structured, human-readable report summarizing its key findings (Supplementary Notes D). It identified a consistent postprandial thermogenic response, with an average temperature rise of 2.19°C (median: 1.10°CC) and a wide range across individuals (-0.11°CC to 15.56°CC). Some participants showed rapid, pronounced spikes within 30 minutes of eating, while others had delayed or muted responses – indicating divergent metabolic phenotypes (Figure 3c,d). These insights were not manually curated or extracted by a human; the agent performed the entire analysis end-to-end and surfaced the results as a concise narrative highlighting patterns that would otherwise being ignored in raw data.

In a parallel workflow, the scientist requested Biomni to analyze 227 nights of wearable-recorded sleep data across 10 participants. Biomni computed averages for duration, efficiency, latency, and sleep stage composition, derived a composite sleep quality score, and conducted chronobiological analyses. The agent delivered a structured summary to the user (Supplementary Notes D, Supplementary Figure 4), including personalized sleep profiles and timing insights, without human post hoc synthesis. Biomni uncovered several novel insights: sleep efficiency consistently peaked mid-week (on Wednesdays) and declined on Sundays, suggesting a potential behavioral pattern tied to pre-Monday stress or weekend-induced disruptions. Another important finding was that consistent sleep timing correlated more strongly with higher sleep quality than total sleep duration, highlighting the critical role of circadian regularity in maintaining restorative sleep.

The scientist then tasked Biomni with analyzing multi-omics data (652 lipidomic, 731 metabolomic, and 1,470 proteomic features), jointly with the CGM data. Biomni conducted cross-omics correlation analysis, applied hierarchical clustering to uncover biologically coherent feature groups, and performed unsupervised PCA to link CGM signals to molecular pathways. It automatically generated interpretable outputs – trajectory plots, heatmaps, boxplots, PCA biplots, and cluster maps – empowering rapid insight generation from complex multimodal datasets (Supplementary Notes D, Supplementary Figure 5). Significant correlations among lipids, metabolites, and proteins revealed tightly interlinked regulatory pathways, underscoring the systems-level nature of metabolic regulation. Notably, several identified biomarkers showed consistent patterns across samples and exhibited high connectivity within correlation networks. Across all cases, the scientist noted that Biomni accelerated the path from messy real-world data to testable hypotheses, supporting applications in sleep optimization, metabolic research, and precision health.

### Biomni automates complex multi-omics analysis to decipher transcriptional regulation of skeletal lineages

To test whether Biomni could generalize to complex omics workflows, a scientist used it to analyze a recently published multi-omics dataset of the developing human skeleton ^27^. This dataset comprises 336,162 single-nucleus RNA-Seq (snRNA) and ATAC-seq (snATAC-Seq), paired with spatial transcriptomics data collected from human embryos between 5-11 weeks post-conception (Figure3e). While the original study emphasized developmental trajectories and disease mechanisms, the scientist was interested in exploring gene regulatory mechanisms across emerging skeletal cell types – a technically demanding task typically requiring extensive bioinformatics support.

The scientist asked Biomni to investigate transcriptional regulation across skeletal lineages using a detailed instruction (Supplementary Notes E). The system autonomously planned and executed a ten-stage analysis pipeline: (1) loading and exploring all datasets, (2) preparing RNA-seq data for analysis, (3) configuring pySCENIC to retrieve motifs, (4) running GRNBoost2 to infer gene regulatory networks, (5) pruning networks using cisTarget, (6) calculating regulon activity with AUCell, (7) extracting accessibility data from ATAC-seq, (8) filtering predicted targets using ATAC-seq accessibility, (9) analyzing activity patterns across cell types, developmental stages, and anatomical regions, and (10) summarizing findings and preparing a report to the scientist. It enabled Biomni to predict transcription factor-target gene links and filter regulons based on motif enrichment and chromatin accessibility correlations (Figure 3f). The full run, completed in just over five hours, handled real-time execution issues (e.g., variable name mismatches) by subsampling and debugging locally. Throughout, Biomni maintained all intermediate outputs – code, figures, and logs – organized in a reproducible folder structure for validation and inspection. The agent summarized all the analysis and generated a report describing the analysis and key findings (Supplementary Notes E).

In its final gene regulatory network (GRN) analysis (Figure 3h), Biomni re-capitulated known regulatory relationships between key osteogenic transcription factors such as RUNX2 and HHIP, confirming how they are regulated by a shared set of anti-osteogenic transcription factors including TWIST1, LMX1B, and ALX4 ^27^. These findings align with author’s report ^27^ about the balanced regulation needed for proper bone formation and suture patency. Furthermore, Biomni also uncovered several unreported TFs, including AUTS2, ZFHX3, and PBX1, showed unexpectedly high regulatory activity across multiple skeletal cell types. Although PBX1 is a well-established skeletal regulator ^28^ and ZFHX3/AUTS2 have only limited or indirect skeletal reports (in mouse ^29^ or zebrafish ^30^), their broad activity here suggests under-appreciated roles across diverse skeletal lineages. Biomni reported that these novel regulators were particularly active in osteoblasts, preosteoblasts, and various chondrocyte populations, suggesting they play important but previously unrecognized roles in the transcriptional control of skeletal cell fate determination during human embryonic development. Finally, Figure 3g-h reveals how Biomni’s visualizations effectively captured both temporal dynamics of regulator activity and cell-type-specific variations in key regulons like RUNX2. This demonstrates how Biomni enables researchers to autonomously perform complex multi-omics analysis and rapidly generate testable hypotheses without specialized programming expertise.

### Biomni designs wet-lab validated experimental protocol for cloning

To evaluate Biomni’s ability to support real-world experimental design, we focused on a core task in molecular biology: cloning. This process is central to countless workflows in research and biotechnology and requires complex reasoning, from designing high-fidelity primers to choosing the right assembly method and validating constructs. While general-purpose LLMs have struggled to perform such tasks due to limited domain knowledge and tool access ^24^, Biomni integrates LLM reasoning with dynamic tool execution, enabling expert-level performance in molecular biology tasks.

To rigorously evaluate this task, we first collaborated with an expert group of gene-editing researchers to design an open-ended cloning benchmark and expert user study (Figure 4a). Our benchmark consisted of 10 realistic, representative cloning tasks covering Golden Gate, Gibson, Gateway, and restriction cloning – each with options including single-fragment vs. pooled assembly. The benchmark also included essential validation steps, such as designing Sanger sequencing primers and analyzing restriction digests. We posed these tasks to four entities: an LLM (Claude 3.7), Biomni, a human trainee (Stanford Biology Master with previous experience in cloning), and a senior human expert (Stanford Genetics PostDoc with 5+ years of cloning experience). Each was asked to generate a complete, end-to-end protocol along with the final cloned plasmid map. Blinded expert reviewers assessed the outputs. Biomni produced protocols and designs that matched the human expert in accuracy and completeness – often providing comparable levels of detail and anticipating the same edge cases. In contrast, the human trainee’s submissions were frequently incomplete or suboptimal, reflecting the experience gap typical in early-stage researchers. Remarkably, Biomni completed all tasks autonomously in a fraction of the time taken by the expert.

**Figure 4:**
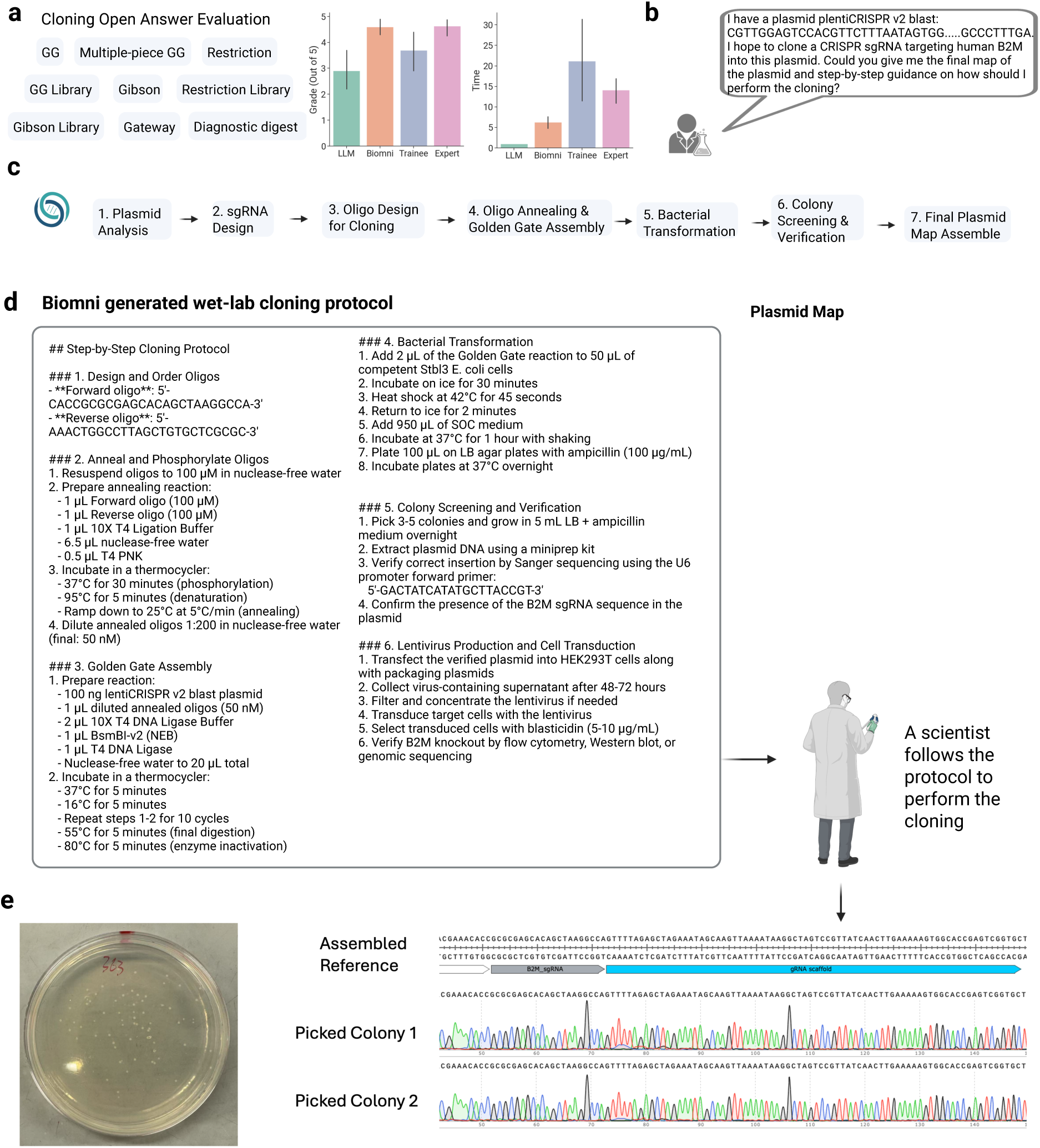
Biomni designs wet-lab experimental protocol. (a) Open-ended cloning benchmark on 10 real cloning scenarios. We compared against base LLM, trainee-level human, and expert-level human scientists. We found that Biomni has similar accuracy as the expert level scientist, and significantly higher accuracy than trainee level, while using much less time. (b) Example of a user request to Biomni for cloning an sgRNA targeting the human B2M gene into the lentiCRISPR v2 Blast plasmid. (c) Biomni’s automated stepwise workflow, including plasmid analysis, sgRNA design, oligo synthesis, Golden Gate assembly, bacterial transformation, colony screening, and final plasmid mapping. (d) Biomni-generated detailed cloning protocol with step-by-step instructions and comprehensive plasmid map, enabling laboratory scientists to execute the experiment autonomously. (e) Validation of Biomni’s cloning protocol through successful colony growth on selection plates, followed by Sanger sequencing confirming perfect alignment of sgRNA insertion in picked colonies, demonstrating Biomni’s robust capability for precise and reliable experimental design.

To further validate Biomni in a real-world setting, a scientist assigned it a practical cloning task: cloning a guide RNA targeting the human B2M gene into the lentiCRISPR v2 Blast construct (Figure 4b). Biomni successfully executed the task through a comprehensive workflow (Figure 4c). First, it analyzed the plasmid structure using annotation and pattern search tools to identify key features necessary for cloning. It then designed three Cas9 sgRNAs targeting B2M using specialized knockout sgRNA design tools. For the cloning process, Biomni generated forward and reverse oligos with BsmBI overhangs to enable directional insertion of the sgRNA sequence. It produced detailed protocols (Figure 4d) for oligo annealing, double-stranded DNA formation, and Golden Gate cloning into the target vector. Biomni also provided complete bacterial transformation instructions, including heat-shock steps and antibiotic selection. For quality control, it designed a U6 promoter sequencing primer to verify sgRNA insertion and simulated the Golden Gate assembly to produce the final plasmid map.

The scientist followed Biomni’s protocol exactly to perform the wet-lab experiment (Figure 4e). Colonies appeared on the plate the next day; two were cultured, miniprepped, and sequenced using the Biomni-designed primers – both showing perfect alignment. This case illustrates how scientists can rely on Biomni to autonomously design complex molecular biology experiments with accuracy comparable to human experts, but in a fraction of the time.

### User-friendly interface to empower scientists to generate biomedical discoveries

To bring the power of Biomni into the hands of every scientist, we built an intuitive graphical interface – available at https://biomni.stanford.edu – to help transform the way researchers interact with biomedical data and tools. This seamless platform enables users to submit natural language queries and receive results powered by the full capabilities of Biomni’s agentic system. Whether designing complex cloning experiments, querying multi-omics databases, or generating hypotheses from wearable data, scientists can now access the intelligence of a general-purpose biomedical AI agent without writing a single line of code. The interface is designed for rapid iteration, real-time feedback, and visual traceability, allowing users to explore intermediate steps, inspect tool usage, and validate results interactively. By closing the gap between biomedical intent and execution, Biomni opens a new era of accessible, automated, and scalable scientific discovery. An example of this interface is shown in Supplementary Figure 17.

## 3 Discussion

Biomni marks a major step forward in biomedical research, demonstrating robust generalization across diverse subfields and laying the groundwork for AI agents as integral collaborators in scientific discovery. Its zero-shot performance across complex tasks – including those in genetics, genomics, microbiology, immunology, pharmacology, and clinical medicine – underscores its potential to boost research productivity, accelerate discovery, and broaden access to advanced biomedical analyses.

By automating complex, labor-intensive workflows, which normally require both expert knowledge and coding skills, Biomni enables researchers to redirect their efforts toward creative hypothesis generation, experimental innovation, and cross-disciplinary collaboration. This shift holds profound implications. In the conext of target and drug discovery for biopharma, Biomni can autonomously prioritize targets, design perturbation screens, or repurpose drugs – offering a path to faster, more cost-effective reasearch. In clinical application settings, its capabilities in gene prioritization and rare disease diagnosis point to more accurate, personalized insights and streamlined diagnostics. For consumer health, Biomni’s integration of wearable data and multi-omics analyses envisions real-time, individualized health monitoring and intervention.

Nonetheless, several limitations remain. While Biomni’s unified environment spans a wide range of biomedical tools and databases, the evaluated tasks represent only a subset of the field, and key domains remain unexplored. In addition, in the action discovery agent, our decision to prioritize the most recent literature makes the agent appear timely, but risks overlooking foundational concepts and techniques that have faded from current discourse despite their enduring relevance. The future versions should encapsulate a larger coverage of publications when defining the environment. Moreover, although Biomni approaches human-level performance in tasks like database querying, sequence analysis, and molecular cloning, it still struggles in areas requiring nuanced clinical judgment, novel experimental reasoning, analytical inventions, or deep biological thinking and synthesis. No system yet captures the full scope of human biomedical expertise. As reflected in our benchmarks, Biomni has not achieved expert-level performance across all task categories. We expect continued improvements as foundation models evolve and the agentic environment expands, as well as thanks to human experts and trainees deploying Biomni to facilitate or augment their work.

These limitations open promising directions for future development. Training biomedical reasoning agents with reinforcement learning could enable continuous self-improvement in planning and execution. Integrating multimodal data – text, images, and structured inputs – may further deepen reasoning capabilities. Equipping Biomni to autonomously discover and incorporate new tools and databases, as well as incorporate more historical methods (which may have high utility but can be easily forgotten by human users), would ensure adaptability and long-term relevance.

Looking ahead, Biomni and its successors could become foundational infrastructure in an AI-powered biomedical ecosystem, working seamlessly with human experts to unlock novel insights into health and disease. This hybrid partnership may radically reshape biomedical research – automating hypothesis generation, scaling discovery pipelines, and enabling medical innovation to proceed at unprecedented speed and scope. General-purpose agents like Biomni could not only accelerate breakthroughs but redefine the future of scientific inquiry itself.

## 4 Methods

### Action Discovery from Literature

100 recent publications from the year 2024 at biorxiv Were collected and analyzed by extracting and parsing their PDF contents. Each paper was processed in chunks, and a specialized prompt guided an LLM through each chunk to explicitly identify and extract three categories of actionable insights: tasks, software, and databases. Specifically for tasks, the LLM was instructed to highlight recurrent tasks requiring specialized implementations within biomedical research workflows.

### Implementing the Biomni Environment

In the initial iteration of environment construction, a conservative and focused approach was adopted for tool curation. Initially, tasks were filtered based on relevance to the primary research interests-drug discovery and clinical biomedicine-retaining fields such as biochemistry, bioengineering, biophysics, cancer biology, cell biology, developmental biology, genetics, genomics, immunology, microbiology, molecular biology, pathology, pharmacology, physiology, synthetic biology, and systems biology. Subsequently, these were narrowed down to approximately 1,900 commonly recurring tasks. These tasks were further manually reviewed to eliminate redundancy and exclude tasks that were trivial or easily implementable through simple code. Selecting highly specialized tasks that require significant domain expertise was emphasized, such as wet-lab protocols and advanced AI models.

Human scientists then collaborated with software engineering agents equipped with web search capabilities to implement each specialized tool. Every tool underwent rigorous validation, requiring a clearly defined test case that it successfully passed. This stringent process culminated in a curated collection of 150 specialized tools. Additionally, essential literature retrieval tools were included, such as PubMed and Google Scholar, with provisions for future iterative expansions.

Each tool was strictly defined using a comprehensive checklist that mandated: (1) a clear and descriptive name, (2) detailed documentation, (3) outputs formatted as detailed research logs optimized for LLM interpretation, (4) the inclusion and successful passing of a specific test case, and (5) specialization criteria-if a task could easily be implemented via brief LLM-generated code (e.g., simple database queries), no specialized tool was created.

Databases were categorized and extensive relational databases accessible via web APIs (e.g., PDB, OpenTargets, ClinVar) were integrated using a unified querying function. This function accepts natural language inputs and leverages an LLM to dynamically parse database schemas and execute corresponding queries. Databases lacking web APIs were downloaded and locally preprocessed into structured pandas DataFrames for seamless accessibility by the agent.

For software integration, recognizing the frequent necessity of concurrently utilizing multiple software tools, a unified containerized environment was constructed, which was pre-installed with a comprehensive suite of relevant software. Additionally, this environment supports the execution of R packages and command-line interface (CLI) tools.

### Biomni-A1

The Biomni agent is a general-purpose biomedical AI agent built upon the CodeAct^31^ framework, designed to systematically solve biomedical tasks by combining LLMs with an interactive coding environment. Given a user query, Biomni begins by prompting the LLM to generate a clear, numbered bullet-list plan detailing the steps needed to tackle the given problem, keeping careful track of progress and adjustments along the way. As the tool, software, and database space is vast, the query task may only use a small set of these resources. To avoid long context, a prompt-based retriever is utilized, powered by a separate LLM, where the agent dynamically selects the most relevant functions, datasets, and software libraries from available resources. During execution, the LLM generates code, executes it in a coding environment (Python, R, or Bash), and returns the resulting observations to inform subsequent reasoning. This iterative approach continues until the agent converges on an accurate, validated solution.

### Q&A Benchmarks

Development and testing sets were created by sampling the LAB-Bench Database Question-Answering and Sequence Question-Answering benchmarks ^24^. Due to resource constraints, each set comprises 12.5% of the complete reference, proportionally distributed across benchmark subtasks, providing a cost-effective and representative assessment of model performance. The development set informed iterative refinements to Biomni’s database integrations and tool implementations, while the test set provided an independent evaluation of generalization capabilities. Accuracy was evaluated by following the LAB-Bench protocol, using multiple-choice answer options with an option for abstention due to insufficient information. Results represent averages across three independent evaluation runs.

For Humanity’s Last Exam (HLE)^23^, a representative sample of questions was selected, spanning fourteen subdisciplines of Biology/Medicine: Genetics, Biology, Ecology, Neuroscience, Biochemistry, Microbiology, Immunology, Molecular Biology, Computational Biology, Biophysics, Bioinformatics, Genomics, and Physiology. From each subdiscipline, up to five questions were sampled (or the maximum number available if fewer than five existed in the category). This sampling approach yielded a final evaluation set of 52 questions that comprehensively assessed Biomni’s performance across the biological sciences. The evaluation was conducted directly without the use of a development set.

### Curating real-world benchmarks

The variant prioritization benchmark was curated from Open Target Genetics^32^ ground truth set, and processed such that given a variant, a negative set of variants is found. The prompt was as follows: “Your task is to identify the most promising variant associated with a given GWAS phenotype for futher examination. From the list, prioritize the top associated variant (matching one of the given variant). GWAS phenotype: {trait} Variants: {variant list}”. Accuracy was used as the metric. The GWAS causal gene detection benchmark utilized a dataset curated from Shringarpure et al^33^, using the original prompt: “Your task is to identify likely causal genes within a locus for a given GWAS phenotype. From the list, provide only the likely causal gene (matching one of the given genes). Identify the causal gene. GWAS phenotype: {trait} Genes in locus: {gene str}”. Accuracy was used as the metric. The perturbation screen design benchmark was curated from Schmidt et al.^34^. The prompt is “Task: Plan a CRISPR screen to{task description}. There are 18,939 possible genes to perturb and only perturb {num genes} genes. For each perturbation, you can measure out {measurement} which will be referred to as the score. Generate {num genes} genes that maximize the perturbation effect. Output format: a list of genes 1. XXX 2.XXX 3.XXX …”. The evaluation metric was the average post-perturbed effect. As the scale differs for the post-perturbed effect, one screen (IL-2) was used. The scRNA-seq annotation benchmark ensured flexibility across diverse data formats (e.g., CellxGene, author-hosted portals), encompassing multiple tissues, species, sequencing technologies, and experimental conditions. Datasets with author-provided annotations (Tier 1 or Tier 2, typically 2≥10 cell types) were prioritized, and 20k-50k cells were subsampled proportionally to their cell type distributions. Automatic evaluation was conducted at the single-cell level using LLMs via *semantic match*, accounting for both naming variations (e.g., fibroblast vs. Fibroblast cells) and hierarchical relations (e.g., CD8+ T cells vs. T cells), judged on-the-fly by LLM agents and later verified by humans. In the microbiome benchmark, both Biomni and human experts independently performed differential abundance analysis on five diverse microbiome datasets, selected to reflect different data types, biological contexts, and analytical challenges. Dataset 1 comes from the MGM 2.0 platform^35^ and includes relative microbial abundance across samples and another with sample labels, ideal for classification tasks^35^. Dataset 2 curated from a well-known Nature study, offers microbial abundance data in mice alongside metadata such as diet and sex, making it valuable for modeling host-microbiome interactions^36^. Dataset 3, developed by Pasolli et al.^37^, combines eight human metagenomic studies with species-level features processed using MetaPhlAn2^37^. Dataset 4 explores microbial communities in drinking water systems, providing an OTU matrix with abundances represented as relative sequence counts. This environmental dataset allows models to be tested beyond host-associated microbiomes^38^. Finally, Dataset 5 is an in-house resource derived from the Human Microbiome Project^39^. Together, these datasets provide a comprehensive foundation for benchmarking AI agents in microbiome analysis across both clinical and environmental domains. Biomni results were compared against those generated by human experts for consistency, accuracy, and efficiency. The drug repurposing benchmark used a dataset from Huang et al.^40^, for the task of identifying the most likely drug from a pre-defined list of drugs for repurposing in a given indication. Evaluation was based on the alignment score with off-label prescription patterns of clinicians from an EHR system. The prompt was “Your task is to identify top 5 drugs that can be potentially repurposed to treat the given disease. From the list, prioritize the drug list with the highest potential (matching the given DrugBank IDs). Disease: {disease} Drugs: {drug list} Output format: a list of drugs with their DrugBank IDs, no drug name, just the IDs: 1. DB00001 2. DB00002 3. DB00003 ..”. The rare disease diagnosis benchmark used the MyGene2 dataset, curated by Alsentzer et al.^41^. The ground truth was expert annotated diagnosis. The prompt was “Task: given a patient’s phenotypes and a list of candidate genes, diagnose the rare disease that the patient has. Phenotypes: {phenotype list} Candidate genes: {candidate genes} Output format: {{’disease name’: XXX, ’OMIM ID’: XXX}}”. The patient gene prioritization benchmark used a dataset curated by Alsentzer et al.^41^. The ground truth was a truly causal gene. The prompt was “Task: Given a patient’s phenotypes and a list of candidate genes, identify the causal gene. Phenotypes: {phenotype list} Candidate genes: {candidate genes} Output format: {{’causal gene’: [gene1]}}”.

### Wearable analysis case study

A wearable case study integrated CGM-derived body temperature data, sleep metrics, and multi-omics datasets from human participants^42^, as follows: CGM Body Temperature Data: For each participant, continuous glucose monitors (CGMs) equipped with temperature sensors recorded skin temperature in high resolution. A total of 485 temperature files were collected, each centered on a presumed meal event. The time window for each file spanned 6 hours total, comprising 2 hours pre-meal and 4 hours post-meal. Sleep Data: Sleep metrics were derived from wrist-worn wearable devices for a subset of 10 participants, covering 227 nights of sleep. Parameters collected included sleep duration, sleep efficiency, sleep latency, sleep stage composition (light, deep, REM), and number of wake episodes. Omics Data: Blood samples were analyzed to generate the following: Lipidomics: 652 lipid features across 147 samples; Metabolomics: 731 metabolite features across 147 samples; Proteomics: 1,470 protein features across 20 samples.

### Multiome analysis case study

The authors’ dataset was directly downloaded and used with no modifications ^27^. The authors’ study generated a multi-omic dataset of human embryonic skeletal development from 5-11 weeks post-conception. The dataset includes snRNA-seq and snATAC-seq data from approximately 336,000 nuclei across five anatomical regions (hip, knee, shoulder joints, calvaria, and skull base). The dataset covers both appendicular (limb) and cranial regions. No additional tools or manual preprocessing were added. As the analytical traces are extensive, more guidance was included in the prompt instruction and two use cases were tested:

*Comparative Analysis.* This analysis focused on how cellular processes differ across anatomical locations and developmental timepoints. Biomni was instructed to characterize the cellular composition across anatomical regions (calvaria, skull base, shoulder, hip, knee) and developmental stages. We prompted Biomni with detailed instructions (Supplementary Section E), including cell type proportion estimates, region-specific population labels, UMAP embeddings, stacked bar plots, a comparison of intramembranous versus endochondral ossification, key transcription factor highlights, and developmental trajectory tracing.

*Gene Regulatory Network Analysis* We asked Biomni to identify transcriptional programs underlying skeletal development. Following a systematic 10-step process, Biomni inferred gene regulatory networks by: (1) loading and exploring all datasets, (2) preparing RNA-seq data for analysis, (3) configuring pySCENIC to retrieve motifs, (4) running GRNBoost2 to infer gene regulatory networks, (5) pruning networks using cisTarget, (6) calculating regulon activity with AUCell, (7) extracting accessibility data from ATAC-seq, (8) filtering predicted targets using ATAC-seq accessibility, (9) analyzing activity patterns across cell types, developmental stages, and anatomical regions, and (10) summarizing findings.

*Manual verification* To evaluate whether the aggregated findings are truly reflected by the data or merely simulated or hallucinated by the LLM, manual (human) verification was conducted following the traces and codes generate by Biomni.

### Wetlab Benchmark Development and Evaluation

A comprehensive benchmark was developed consisting of 20 open-ended cloning questions curated from real-world applications to represent the diversity and complexity of molecular cloning tasks across four major categories: Golden Gate assembly, Gibson assembly, restriction enzyme cloning, and Gateway cloning. Each category included both single-construct and pooled cloning scenarios. Additionally, the benchmark incorporated common validation methods, including diagnostic restriction digestion, Sanger sequencing primer design, and sequence alignment analysis. For establishing baseline performance, three human experts with extensive experience in molecular cloning were recruited. These experts were instructed to complete each task without utilizing language models but were permitted to use standard molecular biology tools, search engines, and publicly available online resources such as plasmid repositories and primer design platforms. The time required for each expert to complete each task was recorded, from initial task understanding to the final protocol and plasmid map generation. In parallel, Biomni and general LLM models were evaluated on identical tasks. Each system was provided with the same task descriptions and required to generate detailed end-to-end experimental protocols and final cloned plasmid maps. For general LLMs, Claude 3.7 was used as one of the most capable publicly-available models at the time of testing, providing it with the same information but without access to specialized molecular biology tools. For evaluation, an independent senior researcher with experience in molecular cloning technologies was recruited and blinded to the source of each protocol (human expert, Biomni, or general LLM). The evaluator assessed each protocol and plasmid map based on two primary criteria: (1) Accuracy: The correctness of the proposed methodology, including appropriate enzyme selection, reaction conditions, primer design parameters, and plasmid construction strategy. (2) Completeness: The thoroughness of the protocol, including all necessary steps, reagents, concentrations, incubation times, and verification methods. Each criterion was scored on a scale of 1-5 according to a detailed rubric (Supplementary Table S31-32). The average scores across all 20 tasks were calculated for each system and human expert to enable direct comparison.

### Wetlab Validation

A practical cloning task was selected for validation: the insertion of a guide RNA targeting the human B2M gene into the lentiCRISPR v2 Blast construct. This task was chosen for its relevance to CRISPR-based gene editing applications and its moderate complexity, involving multiple molecular biology techniques. The experiment was conducted in a standard molecular biology laboratory setting using commercially available reagents and materials. The lentiCRISPR v2 Blast plasmid was obtained from Addgene. All protocols for the experiment were generated entirely by Biomni without modification (Supplementary Notes F), including plasmid analysis, sgRNA design, oligo design with appropriate overhangs, detailed Golden Gate assembly conditions, bacterial transformation parameters, and verification strategies. For validation of the cloning results, standard molecular biology practices were followed, selecting colonies for culture and miniprep, followed by Sanger sequencing using the Biomni-designed primers. Sequence alignment analysis was performed to verify the correct insertion of the sgRNA sequence. The success of the cloning process was determined by the presence of bacterial colonies on selective media and subsequent sequence verification confirming the accurate incorporation of the designed sgRNA construct into the lentiCRISPR v2 Blast backbone.

## Data availability

All data used in Biomni are publicly available at Harvard Dataverse under https://doi.org/10.7910/DVN/CE4ZYG.

## Code availability

Biomni is open-sourced at https://github.com/snap-stanford/biomni. A web-based user interface is available at https://biomni.stanford.edu. Note that the public tool is not for protected health information.

## Acknowledgements

We thank Emily Alsentzer, Andrew Lee, members of Jure Leskovec’s lab, and members of Euan Ashley’s lab, for providing helpful feedbacks. K.H. and J.L. also gratefully acknowledge the support of NSF under Nos. OAC-1835598 (CINES), CCF-1918940 (Expeditions), DMS-2327709 (IHBEM), IIS-2403318 (III); Stanford Data Applications Initiative, Wu Tsai Neurosciences Institute, Stanford Institute for Human-Centered AI, Chan Zuckerberg Initiative, Amazon, Genentech, GSK, Hitachi, SAP, and UCB. K.H. acknowledge the support of Stanford Bio-X fellowship. Research reported in this publication was supported by the National Institute of Neurological Disorders and Stroke of the National Institutes of Health under Award Number U01NS134358. The content is solely the responsibility of the authors and does not necessarily represent the official views of the National Institutes of Health.

## Authors contribution

K.H., Y.R., J.L. conceived the study. K.H. and J.L. supervised the project. K.H. designed and developed the framework. K.H., S.Z., H.W., Y.Q., Y.L. implemented tools and databases. K.H. designed and implemented the generalist agent architecture. K.H. and R.L. designed the action discovery agent. S.Z. performed benchmarks on Q&A tasks. K.H., H.W., Y.L. collected and implemented benchmarks on realistic tasks. X.Z. provided advice on microbiome benchmark. H.W., J.Z., P.H., K.H. performed multi-omics integration case study. Y.L., K.H. performed wearable data analysis case study. Y.Q., J.Z., D.Y., S.Z., Y.L., K.H. performed wetlab case study. K.H., S.M., J.C., M.W., J.B. performed rare disease diagnosis case study. R.L. performed qualitative trace analysis. R.L., L.Q., G.L., provided support for software. K.H., S.Z., H.W., Y.Q., A.R., Y.L. wrote the draft paper. All authors discussed the results and contributed to the final manuscript.

## Competing interests

A.R. and H.W. are employees of Genentech and A.R. has equity in Roche. All other authors declare no competing interests.

