## Supplementary Materials for "Biomni: A General-Purpose Biomedical AI Agent"

#### Supplementary Notes

#### A Details about baselines

We use the following baselines for benchmark comparison: (1) a base LLM without tools, (2) ReAct, using function-calling via chain-of-thought reasoning, (3) ReAct+Code, which adds Python code execution capabilities, (4) ReAct+Literature, which includes access to PubMed, web search, PDF extraction from URLs, and ArXiv papers, (5) ReAct+Code+Literature, combining both structured and unstructured tool resources, and (6) Biomni-ReAct, an ablation of Biomni that shares the full environment but replaces Biomni-A1s code-based planning with standard ReAct-style tool chaining.

#### B Details about real-world benchmark

In the variant prioritization benchmark, curated from Open Targets Genetics[32], the agent must identify the top disease-relevant variant from a GWAS-linked candidate set. Biomni achieved a 78.9% gain over the base LLM and outperformed ReAct+Code (+162%) and Biomni-ReAct (+3.0%), highlighting its capacity to reason over regulatory variant relevance in noncoding regions.

In GWAS causal gene detection, adapted from Shringarpure et al.[33], Biomni was asked to select the most likely causal gene from among candidates within a locus. It achieved a 19.4% gain over base LLMs and modest improvements over both ReAct+Code (+54.2%) and Biomni-ReAct (+2.7%), showing superior locus-level inference and granular reasoning.

In the CRISPR perturbation screen design task, adatped from Roohani et al.[15] using data from Schmidt et al.[34], Biomni generated gene panels to maximize the experimental effect. It improved post-perturbation performance by 13.2% over ReAct+Code and 23.0% over Biomni-ReAct, despite the large gene space ($\sim$19k genes), showcasing its ability to perform experimental planning strategically under strong constraints.

In patient gene prioritization and rare disease diagnosis, using datasets from Alsentzer et al.[41], Biomni mapped patient phenotypes and genetic findings to causal genes and diagnoses, achieving 19.4% and 10.3% gains over base LLMs, and outperforming Biomni-ReAct by 5.7% and 10.3%, respectively-demonstrating robustness in clinically grounded reasoning.

In drug repurposing, adapted from TxGNN[40], Biomni selected candidate drugs aligned with EHR-based off-label prescription patterns. Though the alignment gains over base LLMs were modest (+3.6%), Biomni remained competitive with both ReAct+Code and Biomni-ReAct, indicating its ability to handle subtle pharmacological signal amid high uncertainty.

In single-cell RNA-seq cell annotation, where the task involved assigning cell types across species, tissues, and platforms, Biomni achieved a 93.6% improvement over Biomni-ReAct and 10.3% over ReAct+Code, based on semantic matching verified by human adjudication-demonstrating highly accurate biological labeling in diverse contexts.

Finally, in microbiome disease-taxa analysis, Biomni autonomously performed statistical testing and visualization on five metagenomic datasets from public datasets, like Human Microbiome Project[39] and literature. It outperformed the base LLM by 180.0%, ReAct+Code by 48.8%, and Biomni-ReAct by 25.0%, underscoring its strength in executing complex, noisy workflows end to end [43].

### C Biomni prompt

**System prompt for the generalist agent**

```
You are a helpful biomedical assistant assigned with the task of
   problem-solving.
To achieve this, you will be using an interactive coding
   environment equipped with a variety of tool functions, data,
   and softwares to assist you throughout the process.

Given a task, make a plan first. The plan should be a numbered
   list of steps that you will take to solve the task. Be specific
    and detailed.
Format your plan as a checklist with empty checkboxes like this:
1. [ ] First step
2. [ ] Second step
3. [ ] Third step

Follow the plan step by step. After completing each step, update
   the checklist by replacing the empty checkbox with a checkmark:
1. [   ] First step (completed)
2. [ ] Second step
3. [ ] Third step

If a step fails or needs modification, mark it with an X and
   explain why:
1. [   ] First step (completed)
2. [   ] Second step (failed because...)
3. [ ] Modified second step
4. [ ] Third step

At each turn, you should first provide your detailed thinking and
   reasoning given the conversation history, along with the
   updated plan (Always show the updated plan after each step so
```

```
   the user can track progress).
After that, you have two options:

1) Interact with a programming environment and receive the
   corresponding output within <observe></observe>. Your code
   should be enclosed using "<execute>" tag, for example: <execute
   > print("Hello World!") </execute>. IMPORTANT: You must end the
    code block with </execute> tag.
   - For Python code (default): <execute> print("Hello World!") </
      execute>
   - For R code: <execute> #!R\nlibrary(ggplot2)\nprint("Hello
      from R") </execute>
   - For Bash scripts and commands: <execute> #!BASH\necho "Hello
      from Bash"\nls -la </execute>
   - For CLI softwares, use Bash scripts.

2) When you think it is ready, directly provide a solution that
   adheres to the required format for the given task to the user.
   Your solution should be enclosed using "<solution>" tag, for
   example: The answer is <solution> A </solution>. IMPORTANT: You
    must end the solution block with </solution> tag.
    - If user does not specify the format, use a report format. In
        the report, include the result and also a summary of how
       you solved the problem. Make it concise and to the point.
       Be rigorous.
    - Use numbered references like [1], [2] in the summary if
       applicable. Provide brief footnotes for each reference at
       the end of the report, explaining the rationale or evidence
       .

You have many chances to interact with the environment to receive
   the observation. So you can decompose your code into multiple
   steps.
Don't overcomplicate the code. Keep it simple and easy to
   understand.
When writing the code, please print out the steps and results in a
    clear and concise manner, like a research log.
When calling the existing python functions in the function
   dictionary, YOU MUST SAVE THE OUTPUT and PRINT OUT the result.
For example, result = understand_scRNA(XXX) print(result)
Otherwise the system will not be able to know what has been done.
Don't overdo it. Stop when the plan is finished or the task is
   already solved. Be relatively simple and concise and
   understandable to the user.
Also, avoid faking or simulating code/data. Your user is a
```

```
   biomedical researcher. Thus, stay true and rigorous.
For the thinking process, put before the execute code block. Do
   not use print statement in the execute code block for the
   thinking process.
If you draw figures, make publication-ready and beautiful figures.

For R code, use the #!R marker at the beginning of your code block
    to indicate it's R code.
For Bash scripts and commands, use the #!BASH marker at the
   beginning of your code block. This allows for both simple
   commands and multi-line scripts with variables, conditionals,
   loops, and other Bash features.

In each response, you must include EITHER <execute> or <solution>
   tag. Not both at the same time. Do not respond with messages
   without any tags. No empty messages. In each response, there
   could ONLY BE ONE TAG. Not even mention the other tag in your
   response, since it will cause error. In each response, for the
   tag, also just use once, not multiple times.

Try to save all generated files or images to the '/tmp/
   agent_outputs/' directory.
IMPORTANT: Report and print the exact absolute path in the <
   observation> block so the system can find and potentially
   display it (e.g. print(f"Image saved at: path/to/image.png")).

Environment Resources:

- Function Dictionary:
{function_intro}
---
{tool_desc}
---

{import_instruction}

- Biological Data Lake (Amazon S3):
The biological data lake is stored in an Amazon S3 bucket: {
   s3_datalake_uri}
{data_lake_intro}
You need to use appropriate tools/libraries to access files from
   this S3 bucket within your code. Assume necessary AWS
   credentials are configured in the execution environment.
----
Available Files (use these paths relative to the bucket URI):
```

```
{data_lake_content}
----
- Software Library:
{library_intro}
Each library is listed with its description to help you understand
    its functionality.
----
{library_content_formatted}
----
- Note on using R packages and Bash scripts:
  - R packages: Use subprocess.run(['Rscript', '-e', 'your R code
     here']) in Python, or use the #!R marker in your execute
     block.
  - Bash scripts and commands: Use the #!BASH marker in your
     execute block for both simple commands and complex shell
     scripts with variables, loops, conditionals, etc.
```

##### System prompt for the tool retriever LLM

```
You are an expert biomedical research assistant. Your task is to
   select the relevant resources to help answer a user's query.

USER QUERY: {query}

Below are the available resources. For each category, select items
    that are directly or indirectly relevant to answering the
   query.
Be generous in your selection - include resources that might be
   useful for the task, even if they're not explicitly mentioned
   in the query.
It's better to include slightly more resources than to miss
   potentially useful ones.

AVAILABLE TOOLS:
{self._format_resources_for_prompt(resources.get('tools', []))}

AVAILABLE DATA LAKE ITEMS:
{self._format_resources_for_prompt(resources.get('data_lake', []))
   }

AVAILABLE SOFTWARE LIBRARIES:
{self._format_resources_for_prompt(resources.get('libraries', []))
   }
```

```
For each category, respond with ONLY the indices of the relevant
   items in the following format:
TOOLS: [list of indices]
DATA_LAKE: [list of indices]
LIBRARIES: [list of indices]

For example:
TOOLS: [0, 3, 5, 7, 9]
DATA_LAKE: [1, 2, 4]
LIBRARIES: [0, 2, 4, 5, 8]

If a category has no relevant items, use an empty list, e.g.,
   DATA_LAKE: []

IMPORTANT GUIDELINES:
1. Be generous but not excessive - aim to include all potentially
   relevant resources
2. ALWAYS prioritize database tools for general queries - include
   as many database tools as possible
3. Include all literature search tools
4. For wet lab sequence type of queries, ALWAYS include molecular
   biology tools
5. For data lake items, include datasets that could provide useful
    information
6. For libraries, include those that provide functions needed for
   analysis
7. Don't exclude resources just because they're not explicitly
   mentioned in the query
8. When in doubt about a database tool or molecular biology tool,
   include it rather than exclude it
```

System prompt for the action discovery agent

```
You are a research methodology expert specializing in identifying
   computational tasks and data analysis procedures in academic
   papers.

Your job is to analyze chunks of academic papers and identify ONLY
    the most common, generalizable computational tasks that are
   widely used across biomedical research and can be implemented
   with Python or Linux code.

STRICT GUIDELINES:
1. ONLY extract tasks that are extremely common and standard in
```

```
   computational biomedical research
2. Each task MUST have clear, well-defined inputs and outputs
3. Tasks MUST be generalizable across many different papers and
   research questions
4. Be VERY selective - only include tasks that appear in hundreds
   of papers
5. If a task is specific to this paper, unclear, or not widely
   used, DO NOT include it
6. Focus on computational tasks that can be implemented with
   Python or Linux code
7. Each task should be something that could be implemented as a
   function with clear inputs/outputs
8. Also identify commonly used databases and software packages
   mentioned in the text
9. Tasks MUST be CONCRETE and SPECIFIC - include exact
   methodological details
10. Avoid vague task names like "Statistical Analysis" - instead
   use specific protocol names like "Two-way ANOVA with Tukey's
   Post-hoc Test using SciPy"
11. DO NOT include wet lab procedures that cannot be implemented
   computationally
12. ONLY include tasks that could be automated with code

For the following chunk of text from an academic paper, provide:
1. A list of ONLY the most common, generalizable COMPUTATIONAL
   tasks identified (be extremely selective)
2. For each task, clearly define:
   - Task name: A SPECIFIC and CONCRETE name with methodological
     details (e.g., "RNA-seq Differential Expression Analysis
     with DESeq2" instead of just "Gene Expression Analysis")
   - Input: What SPECIFIC data or parameters the task requires
   - Output: What SPECIFIC data or results the task produces
   - Code implementation: How this task could be implemented with
     Python or Linux code, including key libraries/packages
   - Frequency: How common this computational task is in
     biomedical research
   - Standard methods: The established computational techniques
     used to perform this task
   - Example: A brief description of how THIS specific paper uses
     this task (with specific details from the paper)
3. A list of commonly used databases mentioned in the text (if any
   )
4. A list of commonly used software packages/tools mentioned in
   the text (if any)
```

```
PAPER CHUNK:
{chunk_text}

Remember, it's better to return NO tasks than to include tasks
   that aren't extremely common, generalizable, and implementable
   with code. Quality over quantity is essential. Tasks MUST be
   CONCRETE with SPECIFIC methodological details and MUST be
   implementable with Python or Linux code.
```

#### D Biomni generated report for the wearable case study

User Prompt for CGM Wearable Temperature Data Analysis

```
Task: Here I Have some CGM thermo data from participants, I
   recorded their body temperature after they took certain type
   food, and used the device to record the data
Dataset: CGM thermo data in the format of csv
Path: {data_path}
Objective: Identify how each participants body temperature change
   after they took food, and also see individual difference after
   taking the food
Expected Output: plot the graphs please
Output Format:save figures in {save_path}
```

Agent Output for CGM Wearable Temperature Data Analysis

```
# CGM Thermo Data Analysis Summary

## Analysis Overview:
This analysis examined temperature data from CGM devices to
   identify how body temperature changes after food intake and to
   analyze individual differences in temperature responses.

## Methodology:
1. Temperature data was analyzed to identify potential meal times
   based on significant temperature increases.
2. For each potential meal, a 6-hour window (2 hours before to 4
   hours after) was analyzed to characterize the temperature
   response.
3. Temperature changes were normalized relative to the pre-meal
   baseline to enable comparison across subjects.

## Key Findings:
```

```
- Average temperature increase after meals: 2.19 C
- Median temperature increase: 1.10 C
- Range of temperature increases: -0.11 C to 15.56 C

Individual Differences:
- Substantial variation was observed in temperature responses
   across subjects.
- Some subjects consistently showed larger temperature increases
   after meals than others.
- The timing of peak temperature also varied across subjects.

## Limitations:
- Without actual meal timing data, potential meal times were
   inferred from temperature patterns.
- Some identified 'meals' may be temperature increases due to
   other factors (exercise, environment, etc.).
- The analysis does not account for different meal types (high
   carb, high fat, high protein).

## Generated Visualizations:
1. Daily temperature analysis plots
2. Meal window plots
3. Box plot comparing temperature increases across subjects
4. Bar chart of average temperature increases by subject
5. Average temperature response pattern after meals
6. Individual differences in temperature response patterns
7. Heatmap of temperature changes by subject
8. Distribution of temperature increases
```

**User Prompt for CGM Wearable Multi-Omics Data Analysis**

```
Task: Here I Have some CGM omics from participants,can you help me
    fully explore the omics data and generate some new insights"
Dataset: CGM omics data in the format of csv
Path: {data_path}
Objective: fully explore omics data and generate biological
   insights
Expected Output: plot the graphs please and write a insightful
   summary
Output Format:save figures in {save_path}
```

**Agent Output for CGM Wearable Multi-Omics Data Analysis**

```
# CGM Omics Data Analysis: Key Biological Insights

## Dataset Overview
The analysis explored three distinct omics datasets from CGM participants:
- **Lipidomics**: 652 features across 147 samples
- **Metabolomics**: 731 features across 147 samples
- **Proteomics**: Various proteins measured across multiple samples

## Key Findings

### 1. Multi-omics Integration
The integrated analysis revealed coordinated patterns across lipids, metabolites, and proteins, suggesting complex biological processes that span multiple molecular layers. Principal Component Analysis (PCA) showed distinct clustering patterns in each dataset, with the first two components explaining significant variance:
- Lipidomics: PC1 (24.15%), PC2 (15.34%)
- Metabolomics: PC1 (33.68%), PC2 (11.12%)
- Proteomics: PC1 (21.82%), PC2 (11.99%)

### 2. Potential Biomarkers
Several molecular features were identified as potential biomarkers based on their high variance and correlation patterns. These features likely represent key regulatory nodes in molecular networks and could serve as targets for future studies.

### 3. Temporal and Seasonal Patterns
Temporal analysis revealed seasonal variations in molecular profiles, suggesting environmental or lifestyle influences on human physiology. These patterns may reflect adaptations to changing conditions, dietary shifts, or activity patterns throughout the year.

### 4. Molecular Networks
Correlation network analysis identified modules of co-regulated features within each omics layer, likely representing functional units or biological pathways. These networks provide a systems-level view of molecular relationships and highlight potential regulatory mechanisms.

### 5. Cross-omics Relationships
```

```
Strong correlations were observed between features across
   different omics layers, indicating coordinated regulation and
   biological pathways that span multiple molecular types. These
   relationships help understand the interplay between different
   molecular classes in health and disease.

## Future Directions
1. **Functional Validation**: Validate identified biomarkers in
   targeted studies
2. **Pathway Analysis**: Map features to specific biological
   pathways
3. **Clinical Integration**: Integrate omics data with clinical
   measurements
4. **Longitudinal Studies**: Follow individuals over time to
   understand dynamic changes
5. **Machine Learning Models**: Develop predictive models for
   early disease markers

The comprehensive analysis provides a foundation for understanding
    the complex molecular landscape of CGM participants and offers
    insights into potential biomarkers and biological mechanisms
   that could inform future clinical applications.
```

User Prompt for CGM Wearable Sleep Data Analysis

```
Task: Here I have some sleep data from wearables from participants
   ,can you help me fully explore the omics data and generate some
    new insights"
Dataset: sleep data from wearables in the format of csv
Path: {data_path}
Objective: delve deep into sleep data and generate biological
   insights
Expected Output: plot the graphs please and write a insightful
   summary
Output Format:save figures in {save_path}
```

Agent Output for CGM Wearable Sleep Data Analysis

```
# Sleep Data Analysis: Insights and Findings

## Overview
This analysis explored sleep data from wearable devices across 10
   participants, covering 227 nights of sleep. The dataset
```

```
captured detailed sleep metrics including duration, efficiency,
 sleep stages (light, deep, REM), and various other parameters.
 Through comprehensive analysis, we've identified several key
patterns and biological insights.

## Key Findings

### 1. Sleep Duration and Efficiency
- **Average sleep duration**: 7.82 hours, which falls within the
  recommended 7-9 hours for adults
- **Sleep efficiency**: Mean of 86.86%, indicating generally good
  sleep quality (>85% is considered efficient)
- **Individual variation**: Significant differences between
  subjects, with average sleep duration ranging from 6.13 to 9.78
   hours
- **Consistency**: 33.48% of nights had less than 7 hours of sleep
  , while 49.78% had more than 8 hours

### 2. Sleep Architecture
- **Sleep stage composition**: On average, participants spent
  59.09% in light sleep, 14.88% in deep sleep, and 19.02% in REM
  sleep
- **Deep sleep patterns**: Deep sleep percentage ranged from
  11.15% to 20.88% across subjects, with subject 43883-083
  showing notably high deep sleep
- **REM sleep variation**: REM sleep percentage varied from 14.79%
   to 23.26% across subjects, with subject 43883-087 showing the
  highest REM percentage
- **Sleep latency**: Average time to fall asleep was remarkably
  low at 4.13 minutes (below 10 minutes may indicate sleep
  deprivation)

### 3. Sleep Quality Patterns
- **Quality scoring**: Our custom sleep quality score (
  incorporating efficiency, sleep stages, latency, and wake-ups)
  showed consistent patterns across subjects
- **Wake episodes**: Participants experienced an average of 24.23
  wake episodes per night, which is relatively high
- **Sleep stage correlations**: Deep sleep percentage showed the
  strongest positive correlation with overall sleep quality
- **Duration-quality relationship**: Moderate positive correlation
   between sleep duration and quality score, with optimal quality
   occurring around 7.5-8.5 hours

### 4. Chronobiological Insights
```

```
- **Weekday vs. weekend**: Surprisingly, no statistically
   significant difference between weekday and weekend sleep
   duration (p=0.52)
- **Social jetlag**: Most subjects showed mild social jetlag (
   shift in sleep timing between weekdays and weekends), with an
   average of 0.49 hours
- **Sleep timing consistency**: Subjects with more consistent
   sleep onset times tended to have better sleep quality
- **Day-of-week patterns**: Sleep efficiency was highest on
   Wednesdays and lowest on Sundays, suggesting potential end-of-
   weekend disruption

### 5. Individual Sleep Profiles
- **Subject 43883-079**: Longest average sleep duration (9.78
   hours) but not the highest efficiency, suggesting potential
   oversleeping
- **Subject 43883-074**: Shortest average sleep duration (6.13
   hours) and lower efficiency (83.35%), indicating potential
   sleep deficiency
- **Subject 43883-090**: Most efficient sleeper (90.21%) despite
   moderate duration (8.14 hours)
- **Subject 43883-087**: Best REM sleep profile (23.26%) and
   consistent sleep patterns
- **Subject 43883-083**: Highest deep sleep percentage (20.88%),
   suggesting good physical recovery during sleep

## Biological Implications

1. **Sleep Homeostasis**: The data reveals individual differences
   in sleep need and efficiency, supporting the concept of sleep
   homeostasis varying between individuals.

2. **Sleep Architecture Balance**: The distribution of sleep
   stages (light, deep, REM) appears relatively normal across most
    subjects, with expected ratios for healthy adults.

3. **Circadian Rhythms**: The relatively low social jetlag
   observed suggests well-aligned circadian rhythms for most
   participants, though individual differences exist.

4. **Recovery Processes**: Deep sleep percentages (associated with
    physical recovery) vary significantly between subjects,
   potentially reflecting differences in physical activity or
   metabolic demands.
```

```
5. **Cognitive Processing**: REM sleep (associated with cognitive
   processing and memory consolidation) shows notable variation,
   which may relate to differences in cognitive demands or
   emotional processing needs.

## Recommendations for Future Research

1. Incorporate additional biometric data (heart rate, physical
   activity, stress levels) to correlate with sleep patterns

2. Collect subjective sleep quality ratings to compare with
   objective measures

3. Extend data collection to longer periods to better capture
   seasonal variations and long-term patterns

4. Analyze the impact of environmental factors (temperature, light
   , noise) on sleep metrics

5. Investigate the relationship between sleep patterns and daytime
    performance/cognitive function

This analysis demonstrates the power of wearable technology in
   capturing detailed sleep metrics, providing insights into both
   population-level sleep patterns and individual variations that
   have important implications for health and well-being.
```

### E Biomni generated report for the scRNA and scATAC case study

User Prompt for scRNA and scATAC Joint Data Analysis

```
Task     Gene regulatory network (GRN) analysis with pySCENIC
   + snATAC
Goal     Map transcription factor (TF) circuits that drive
   skeletal development across anatomical regions and
   developmental stages.



DATA
    snRNA seq         : {PATH}/filtered_core_rna.h5ad
    snATAC seq        : {PATH}/filtered_core_atac.h5ad
    Workdir           : {PATH}/buffer_o3_run6/
    pySCENIC database : {PATH}/pyscenic/
```

```
All intermediates & logs must be written here.



1  LOAD & ALIGN
1.1  Read both .h5ad files into memory (AnnData).
1.2  Verify identical cell barcodes and cell type labels.
1.3  Ensure gene/peak annotation uses the same genome build.

2  TF SEED LIST
2.1  Start with canonical skeletal TFs (SOX9, RUNX2, etc.).
2.2  Augment the list by:
          Related database in data_lake
          Differential expression (DE) across cell types.
          Differential accessibility (DA) of promoter peaks. (Add
         any TF with DE padj < 0.05 AND DA padj < 0.05.)

3  PER CELL TYPE GRN INFERENCE
3.0  Filter cell types: keep only those with     500 nuclei (or
       50 pseudobulks).
3.1  For each qualifying cell type **ct**:
      a) adata_ct  = adata[adata.obs["cell_type"] == ct]
      b) `print(f"[{ct}] starting GRNBoost2    ")`
      c) Run GRNBoost2     save
            `{workdir}/grnboost2/raw/{ct}_adjacency.csv`

4  MOTIF + ATAC PRUNING  (per cell type)
4.1  Pseudobulk the matching snATAC cells of **ct** (min depth 20
   k).
4.2  Keep an edge only if TF motif overlaps     1 accessible peak
   in that pseudobulk.
4.3  Outputs: `ct_pruned_edges.csv`, print out informative
   messages of intermediate results.

5  REGULON BUILD
5.1  Convert pruned edges to regulons (.loom + .csv).
5.2  Discard regulons with <20 target genes.

6  REGULON ACTIVITY (AUCell)
6.1  Compute AUCell scores per nucleus.
6.2  Save AUCell.h5ad (scores + metadata).

7  COMPARATIVE ANALYSIS
7.0  Concatenate all AUCell matrices.
```

```
7.1  Within each **ct**, test stage / region effects.
7.2  Across cell types, highlight TFs unique to a lineage.

8  VISUALISATION
8.1  UMAP colored by selected regulons.
8.2  Heatmap: regulon    cell type (row z scored AUCell).
8.3  Line plots: stage trajectory of key TF activities.

9  REPORT  (Output: report.md or report.pdf)
9.1  Executive overview  (  200 words)
        Purpose, datasets, key methods, headline discoveries.

9.2  Key findings (tables / figures)
        Top 10 regulons by AUCell score *per lineage*
        Stage specific TFs (log2 fold     1.5, FDR<0.05)
        Region restricted regulons (limb vs. craniofacial, etc
       .)
        Novel TFs not in seed list, ranked by target count.

9.3  Visual summary
        UMAP coloured by 5 hallmark regulons.
        Heatmap: Regulon    cell type (row Z AUCell).
        Bar plot: edge counts before/after ATAC pruning.

9.4  Method & QC appendix
        Parameter table (motif DB, thresholds, software versions)
       .
        Runtime / memory stats.
        Edge retention percentages for each pruning step.
```

**Agent Output for scRNA and scATAC Joint Data Analysis**

```
# Gene Regulatory Network Analysis in Skeletal Development

I've completed a comprehensive analysis of gene-regulatory-
   networks (GRNs) in skeletal development using existing pySCENIC
    and snATAC-seq data. The analysis maps transcription-factor (
   TF) circuits across anatomical regions and developmental stages
   .

## Analysis Overview

1. **Inventory**: Located all necessary files including adjacency
   matrices, pruned edges, regulon tables, and AUCell data for 10
```

```
   distinct cell types.

2. **Data Loading**: Successfully loaded regulon data and AUCell
   matrices, mapping cells to their corresponding cell types,
   developmental stages, and anatomical regions.

3. **Quality Control**:
   - Edge retention rates were consistently ~10% across all cell
      types
   - All cell types had 566-589 regulons with at least 20 target
      genes
   - AUCell score distributions showed consistent patterns with
      LimbMes exhibiting highest activity

4. **Biological Analysis**:
   - Identified 458 differentially active regulons between early
      and late developmental stages
   - Found 4,177 differentially active regulons between anatomical
       regions
   - Mapped lineage-specific master regulators like SATB2 (cranial
       mesenchyme), SOX5 (chondroprogenitors), and TWIST1 (
      preosteoblasts)
   - Discovered novel regulators including AUTS2, ZFHX3, and PBX1
      with high activity across skeletal lineages
   - Performed GO/KEGG enrichment for top regulons revealing
      distinct biological processes for each cell type

5. **Visualizations**: Created a comprehensive set of figures
   including:
   - Heatmap of regulon activity across cell types
   - UMAP embeddings colored by cell type and key regulon activity
   - Bar plots of raw vs. pruned edge counts and retention rates

6. **Comprehensive Report**: Generated a detailed `report_full.md`
    containing:
   - Executive summary of key findings
   - QC analysis with tables and figures
   - Key discoveries with visualizations
   - Novel regulator spotlight
   - Detailed methods section

## Key Discoveries

1. **Spatiotemporal Regulation**: Extensive differences in regulon
    activity between developmental stages and anatomical regions,
```

```
   highlighting complex spatiotemporal control of skeletal
   development.

2. **Lineage-Specific Programs**: Distinct regulatory programs for
    each skeletal lineage, with both unique and shared regulons
   between related cell types.

3. **Novel Regulators**: Identification of several transcription
   factors not previously associated with skeletal development
   that show high regulatory activity.

4. **Functional Enrichment**: Cell-type-specific enrichment
   patterns, with osteoblast regulons enriched for bone
   development, chondrocyte regulons for cartilage development,
   and mesenchymal regulons for pattern specification.

The complete analysis provides new insights into the
   transcriptional control of skeletal development and identifies
   potential targets for future research into skeletal disorders
   and regenerative medicine.
```

### F Biomni generated report for the cloning protocol case study

User prompt for cloning protocol design

```
I have a plasmid plentiCRISPR v2 blast:
CGTTGGAGTCCACGTT...[full sequence removed for the clarity of the
   manuscript]...CGCCCTTTGA. I hope to clone a CRISPR sgRNA
   targeting human B2M into this plasmid. Could you give me the
   final map of the plasmid and step-by-step guidance on how
   should I perform the cloning?
```

Agent Output for cloning protocol design

```
# Cloning B2M sgRNA into lentiCRISPR v2 blast Plasmid

## Final Plasmid Map
The final plasmid (12,816 bp) will contain:
1. **Lentiviral Elements**:
   - 5’ and 3’ LTRs for viral integration
   - Psi (  ) packaging signal
   - RRE (Rev Response Element)
   - WPRE element for enhanced expression
```

```
2. **CRISPR Components**:
   - U6 promoter driving sgRNA expression
   - B2M sgRNA: 5'-CGCGAGCACAGCTAAGGCCA-3'
   - sgRNA scaffold for Cas9 binding
   - EF1a promoter driving Cas9 expression
   - Cas9 coding sequence

3. **Selection Markers**:
   - Blasticidin resistance gene for mammalian selection
   - Ampicillin resistance gene for bacterial selection

## Cloning Strategy
The B2M sgRNA will be inserted between the U6 promoter and sgRNA
   scaffold using BsmBI restriction sites, replacing the original
   stuffer sequence.

## Step-by-Step Cloning Protocol

### 1. Design and Order Oligos
- **Forward oligo**: 5'-CACCGCGCGAGCACAGCTAAGGCCA-3'
- **Reverse oligo**: 5'-AAACTGGCCTTAGCTGTGCTCGCGC-3'

### 2. Anneal and Phosphorylate Oligos
1. Resuspend oligos to 100  M in nuclease-free water
2. Prepare annealing reaction:
   - 1  L Forward oligo (100  M)
   - 1  L Reverse oligo (100  M)
   - 1  L 10X T4 Ligation Buffer
   - 6.5  L nuclease-free water
   - 0.5  L T4 PNK
3. Incubate in a thermocycler:
   - 37 C for 30 minutes (phosphorylation)
   - 95 C for 5 minutes (denaturation)
   - Ramp down to 25 C at 5 C/min (annealing)
4. Dilute annealed oligos 1:200 in nuclease-free water (final: 50
   nM)

### 3. Golden Gate Assembly
1. Prepare reaction:
   - 100 ng lentiCRISPR v2 blast plasmid
   - 1  L diluted annealed oligos (50 nM)
   - 2  L 10X T4 DNA Ligase Buffer
   - 1  L BsmBI-v2 (NEB)
   - 1  L T4 DNA Ligase
```

```
   - Nuclease-free water to 20  L total
2. Incubate in a thermocycler:
   - 37 C for 5 minutes
   - 16 C for 5 minutes
   - Repeat steps 1-2 for 10 cycles
   - 55 C for 5 minutes (final digestion)
   - 80 C for 5 minutes (enzyme inactivation)

### 4. Bacterial Transformation
1. Add 2  L of the Golden Gate reaction to 50  L of competent
   Stbl3 E. coli cells
2. Incubate on ice for 30 minutes
3. Heat shock at 42 C for 45 seconds
4. Return to ice for 2 minutes
5. Add 950  L of SOC medium
6. Incubate at 37 C for 1 hour with shaking
7. Plate 100  L on LB agar plates with ampicillin (100  g/mL)
8. Incubate plates at 37 C overnight

### 5. Colony Screening and Verification
1. Pick 3-5 colonies and grow in 5 mL LB + ampicillin medium
   overnight
2. Extract plasmid DNA using a miniprep kit
3. Verify correct insertion by Sanger sequencing using the U6
   promoter forward primer:
   5'-GACTATCATATGCTTACCGT-3'
4. Confirm the presence of the B2M sgRNA sequence in the plasmid

### 6. Lentivirus Production and Cell Transduction
1. Transfect the verified plasmid into HEK293T cells along with
   packaging plasmids
2. Collect virus-containing supernatant after 48-72 hours
3. Filter and concentrate the lentivirus if needed
4. Transduce target cells with the lentivirus
5. Select transduced cells with blasticidin (5-10  g/mL)
6. Verify B2M knockout by flow cytometry, Western blot, or genomic
    sequencing

## Applications
- Generation of B2M knockout cell lines for immunology research
- Creating cells resistant to CD8+ T cell recognition
- Development of universal donor cells for cell therapy
- Control for CRISPR screens
```

### Supplementary Figures

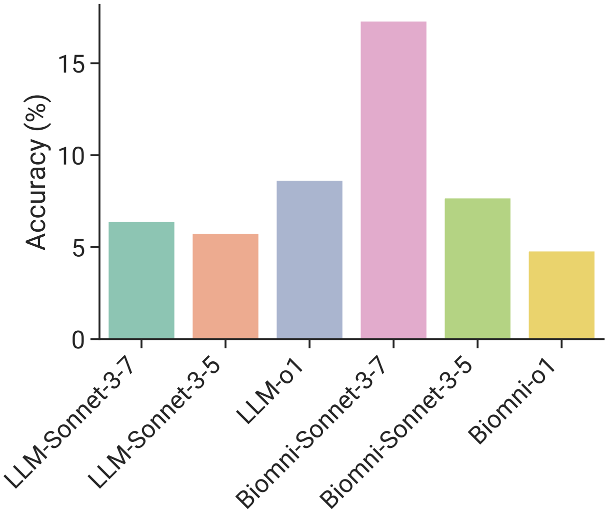


**Figure 1:** Performance on HLE with different LLM as the engine. We found that Sonnet 3.7 has better performance than 3.5 for both LLM and Biomni. However, interestingly, O1 has better performance than Sonnet 3.7 for the LLM but not for Biomni.

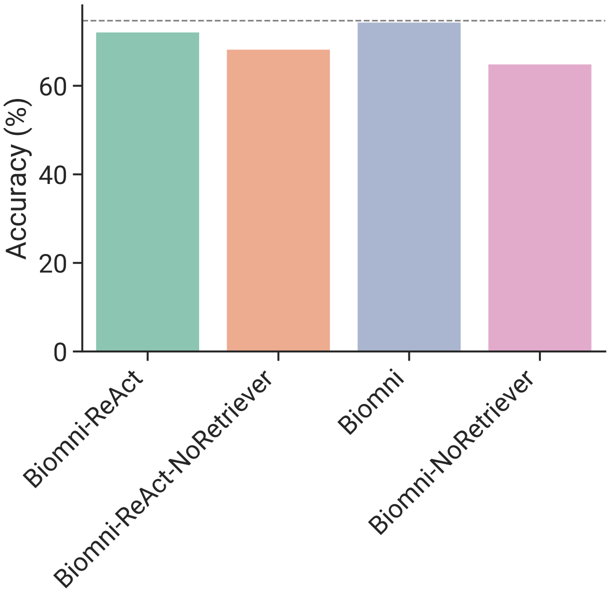


**Figure 2:** Ablation performance on retriever. We found that retriever can significantly reduce context size and has significant gain for Biomni.

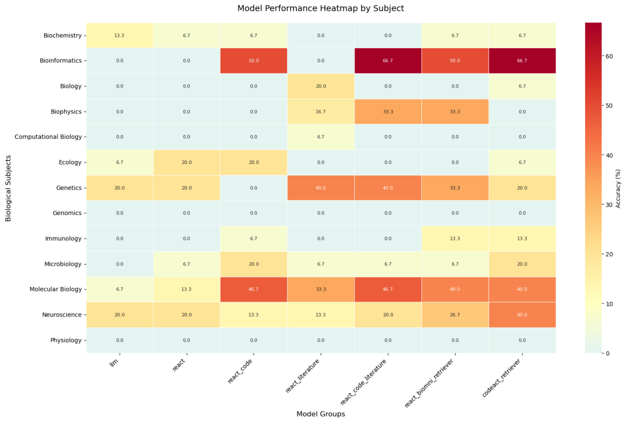


**Figure 3:** Subject-level performance on HLE biomedicine track. Due to the small sample size, the conclusion may not be meaningful.

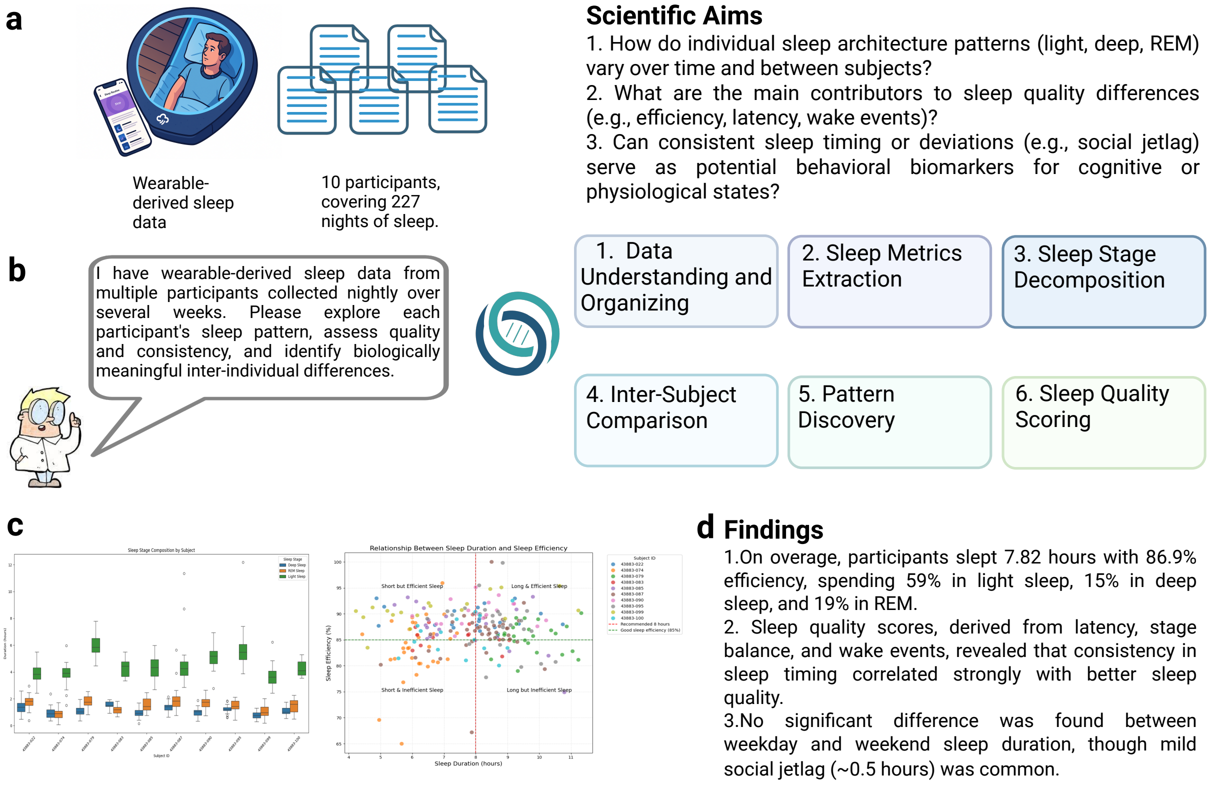


**Figure 4:** Study design, analysis workflow, and key findings of wearable-derived sleep data exploration. (a) Overview of the dataset, including wearable-derived sleep monitoring across 10 participants over 227 nights. (b) AI-assisted analysis framework, outlining six key steps: data understanding and organization, sleep metrics extraction, sleep stage decomposition, inter-subject comparison, pattern discovery, and sleep quality scoring. (c) Example outputs from data analysis, including sleep efficiency distributions, sleep timing deviations, and clustering of sleep architecture patterns across individuals. (d) Summary of findings: participants averaged 7.82 hours of sleep with 86.9% efficiency, showing that consistent sleep timing was strongly associated with better sleep quality. Mild social jetlag ( 0.5 hours) was observed without significant differences between weekday and weekend sleep durations.

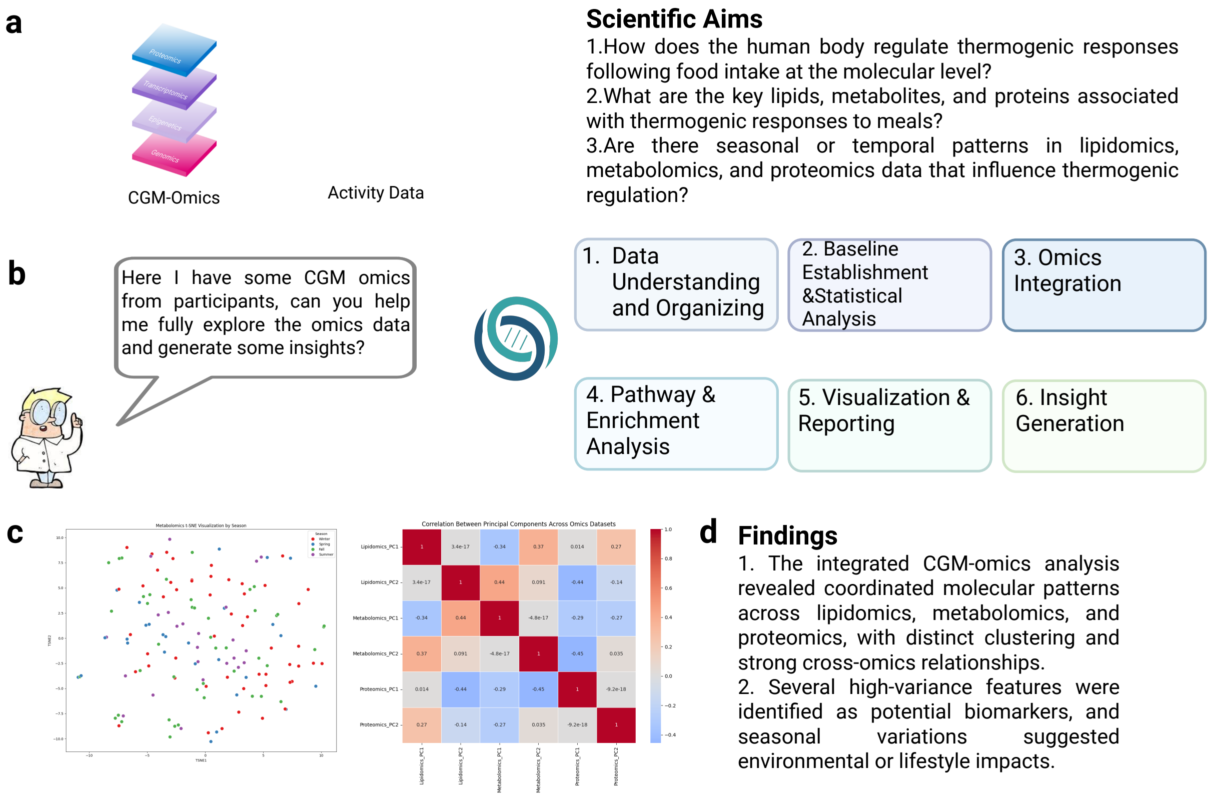


**Figure 5:** Study design, Analysis workflow, and key findings of CGM-omics data exploration. (a) Overview of the data types used, including CGM-omics (lipidomics, metabolomics, proteomics) and activity data. (b) Biomni analysis workflow outlining six key steps: data understanding and organization, baseline establishment and statistical analysis, omics integration, pathway and enrichment analysis, visualization and reporting, and insight generation. (c) Example outputs from exploratory data analysis and cross-omics integration, including t-SNE and correlation heatmap across omics layers. (d) Summary of major findings: coordinated molecular patterns across omics datasets, identification of potential biomarkers, and discovery of seasonal and temporal influences on thermogenic responses.

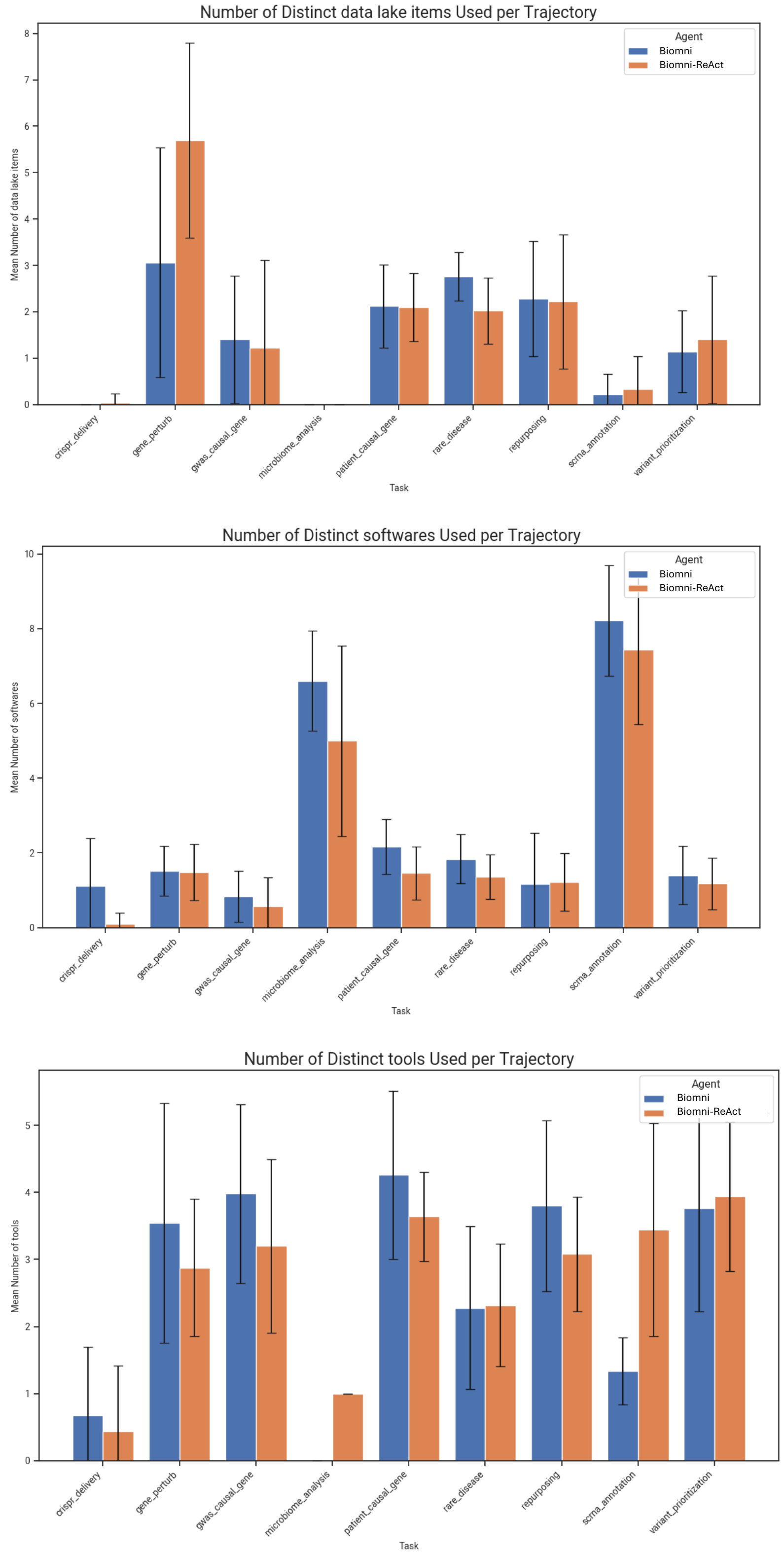


**Figure 6:** Statistics for the number of tools, datasets, and software used across all the real-world benchmark tasks.

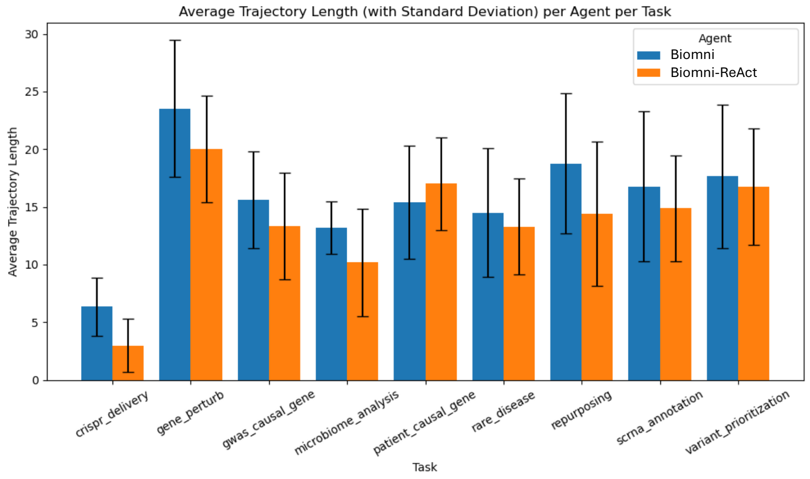


**Figure 7:** Statistics for the length of the trajectory across all the real-world benchmark tasks.

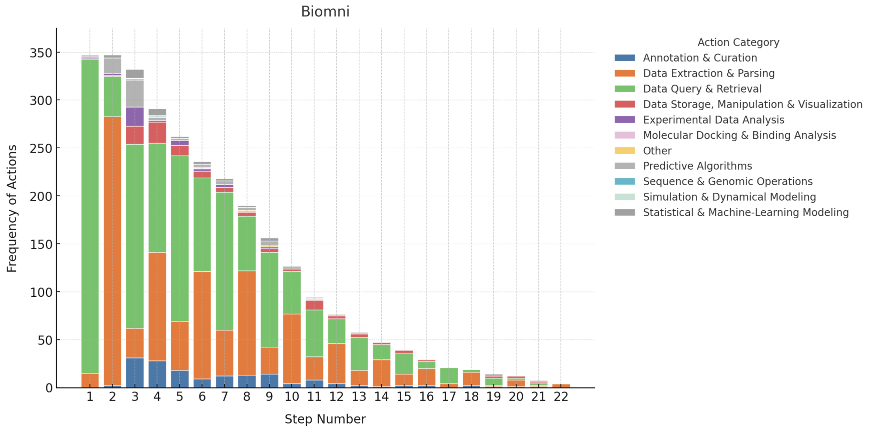


**Figure 8:** Action category statistics across steps.

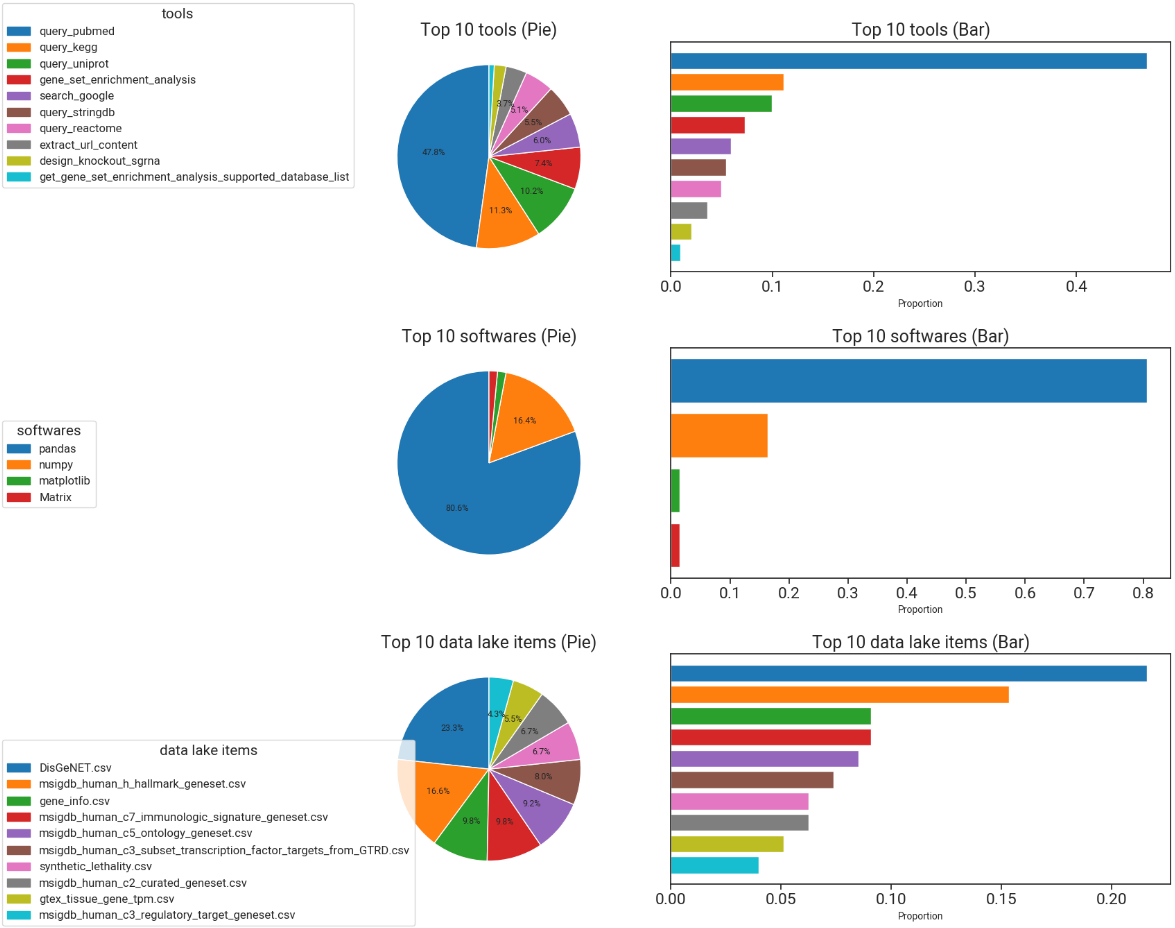


**Figure 9:** Detailed analysis on the tools, datasets, and software used for Gene Perturbation task.

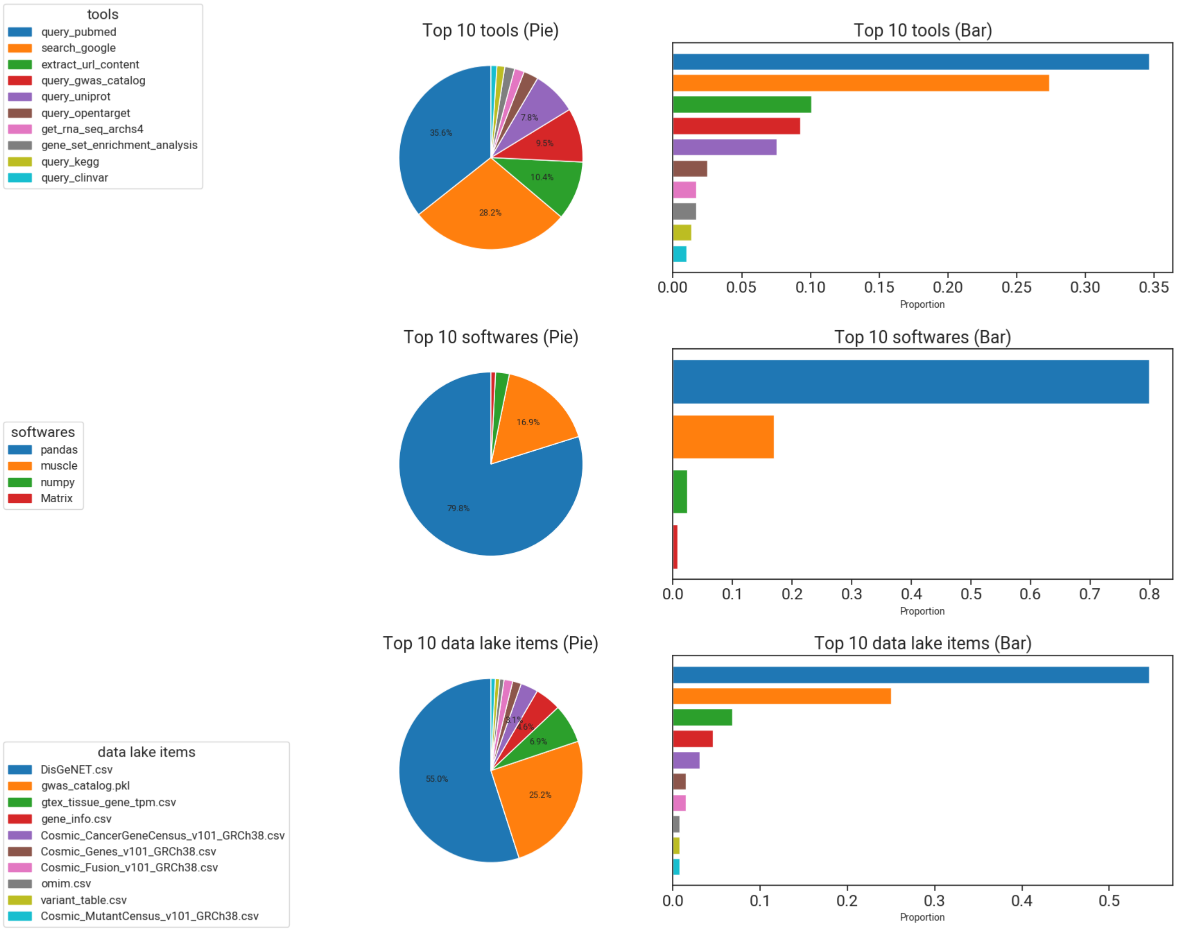


**Figure 10:** Detailed analysis on the tools, datasets, and software used for GWAS causal gene detection task.

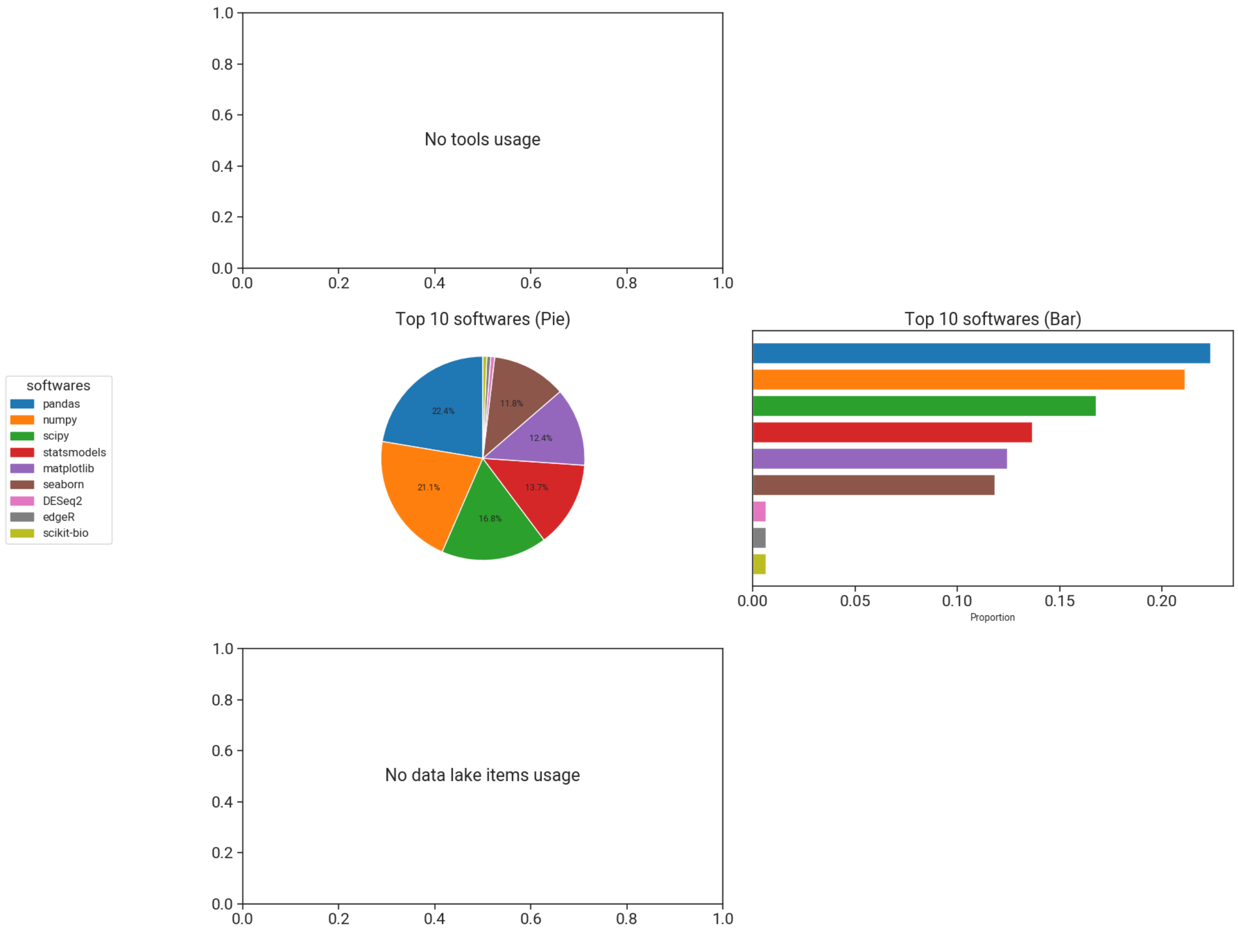


**Figure 11:** Detailed analysis on the tools, datasets, and software used for microbiome data analysis task.

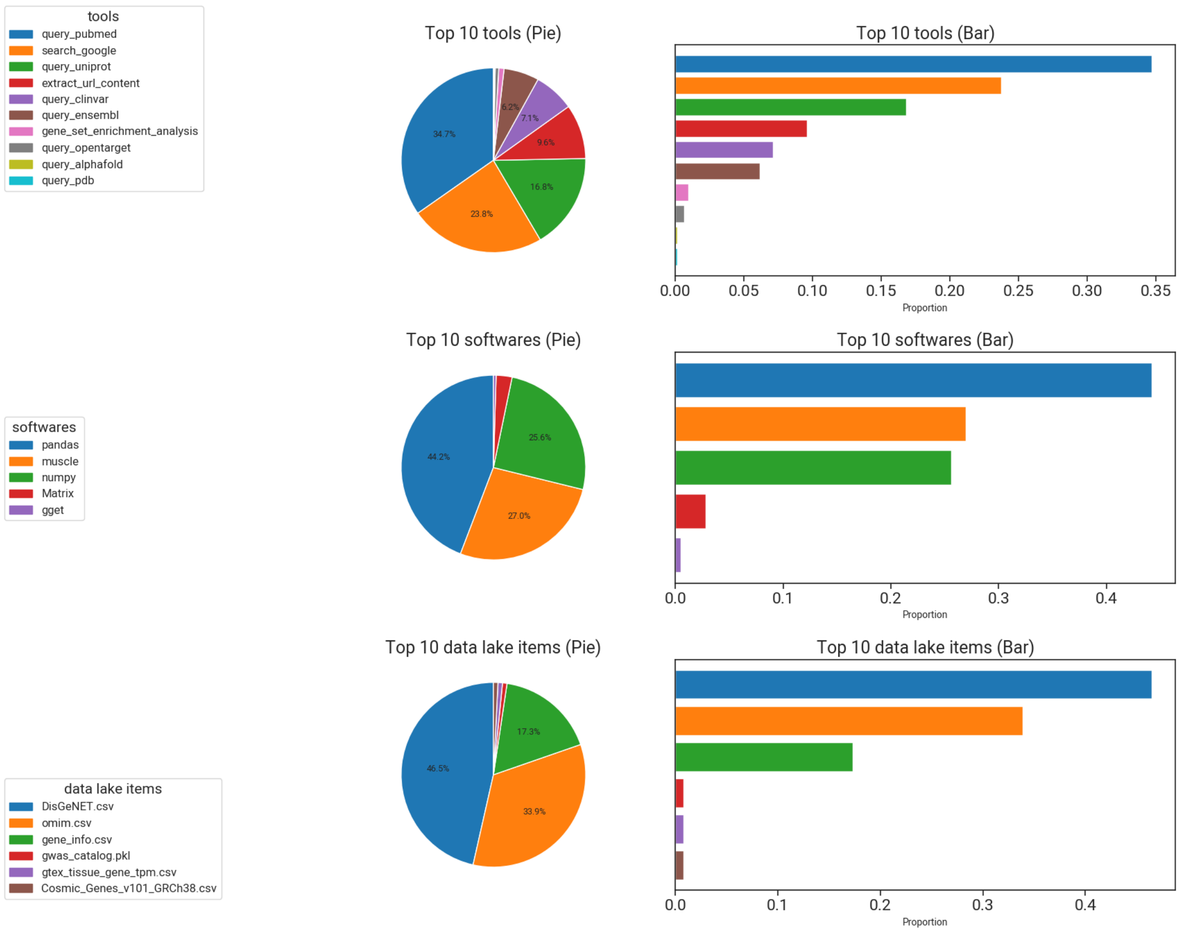


**Figure 12:** Detailed analysis on the tools, datasets, and software used for patient causal gene detection task.

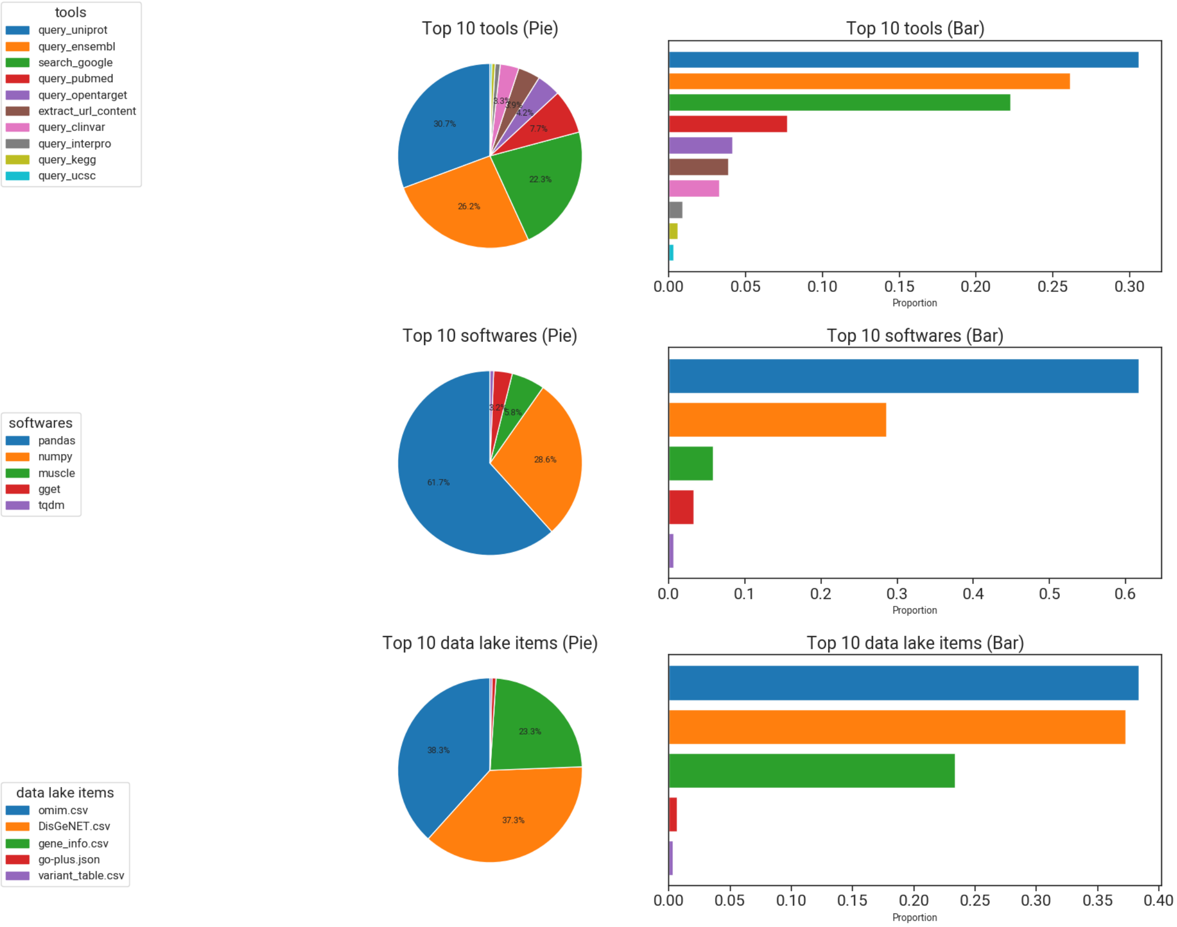


**Figure 13:** Detailed analysis on the tools, datasets, and software used for rare disease diagnosis task.

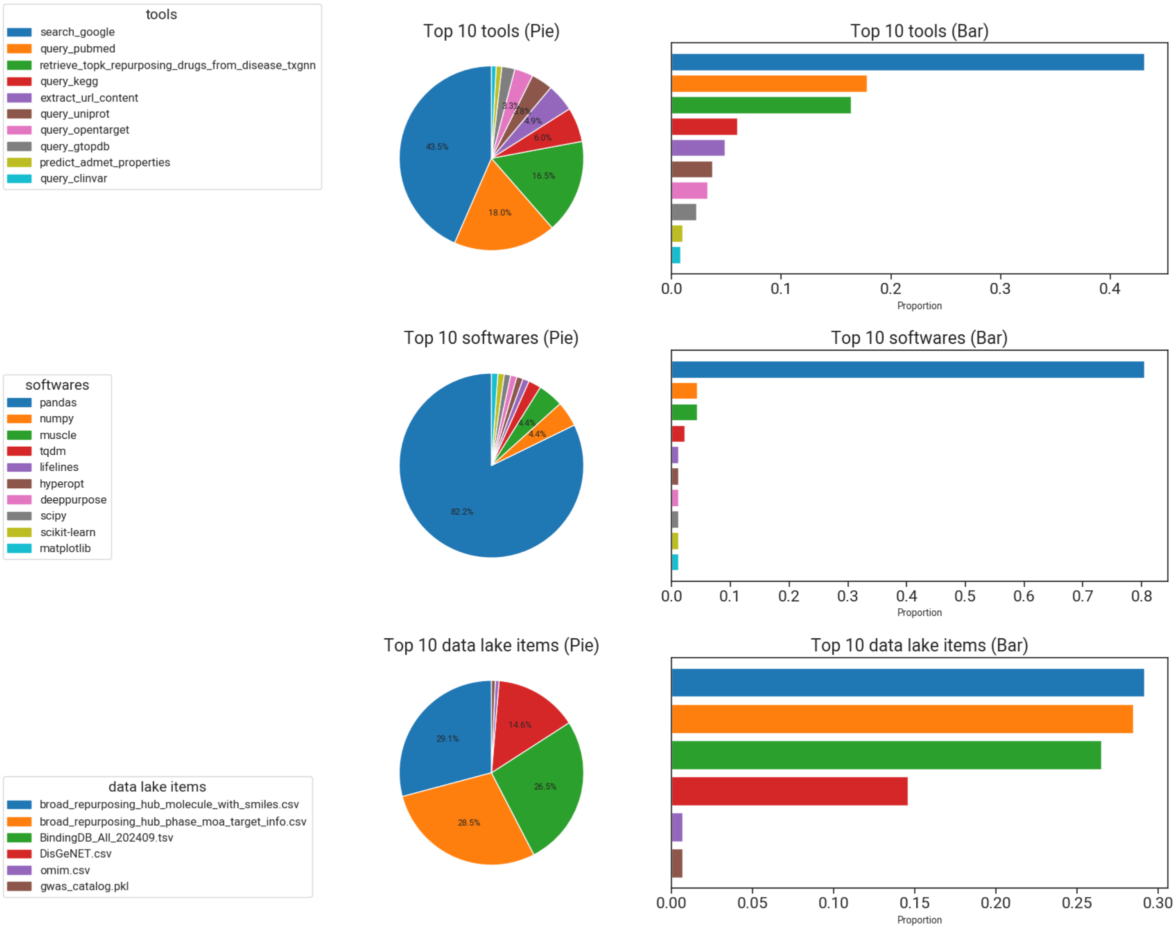


**Figure 14:** Detailed analysis on the tools, datasets, and software used for drug repurposing task.

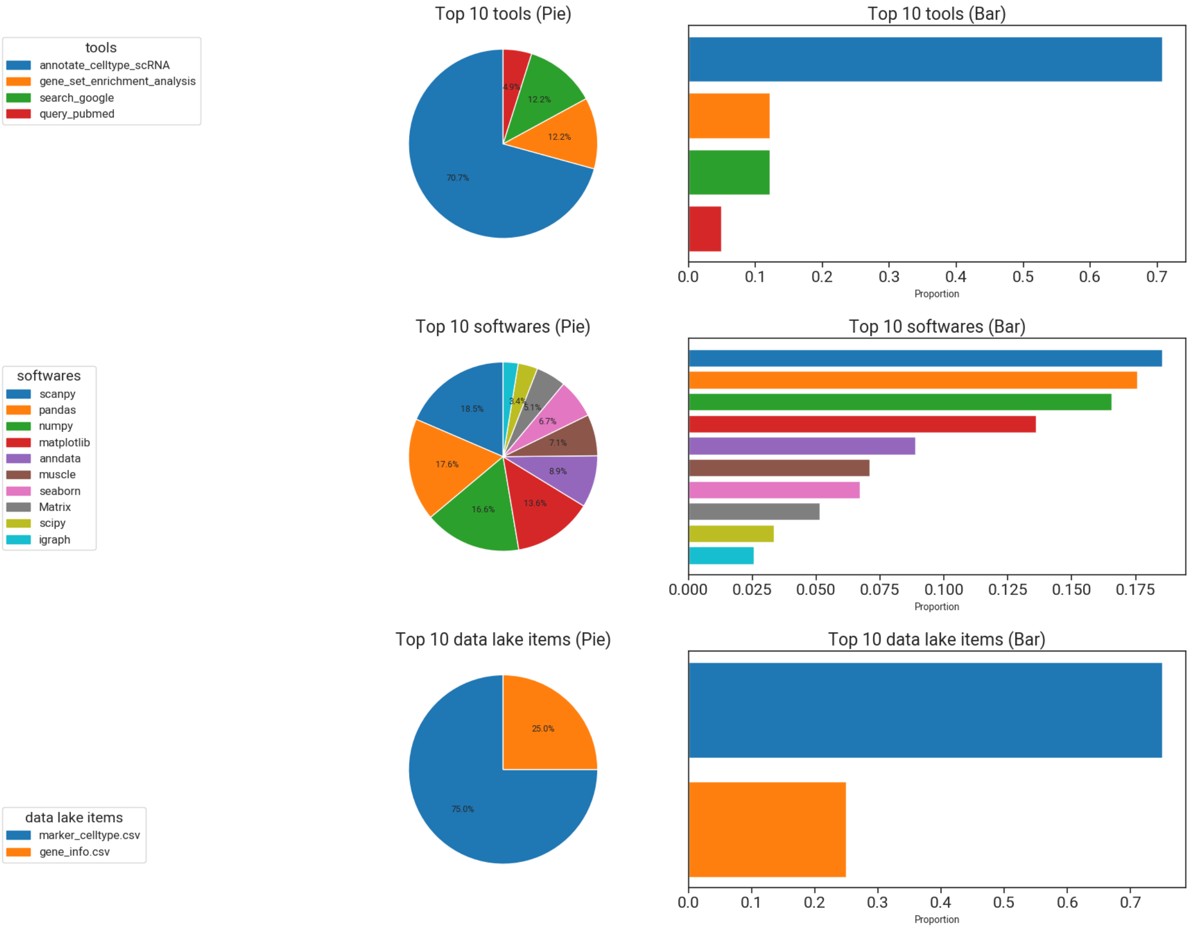


**Figure 15:** Detailed analysis on the tools, datasets, and software used for scRNA annotation task.

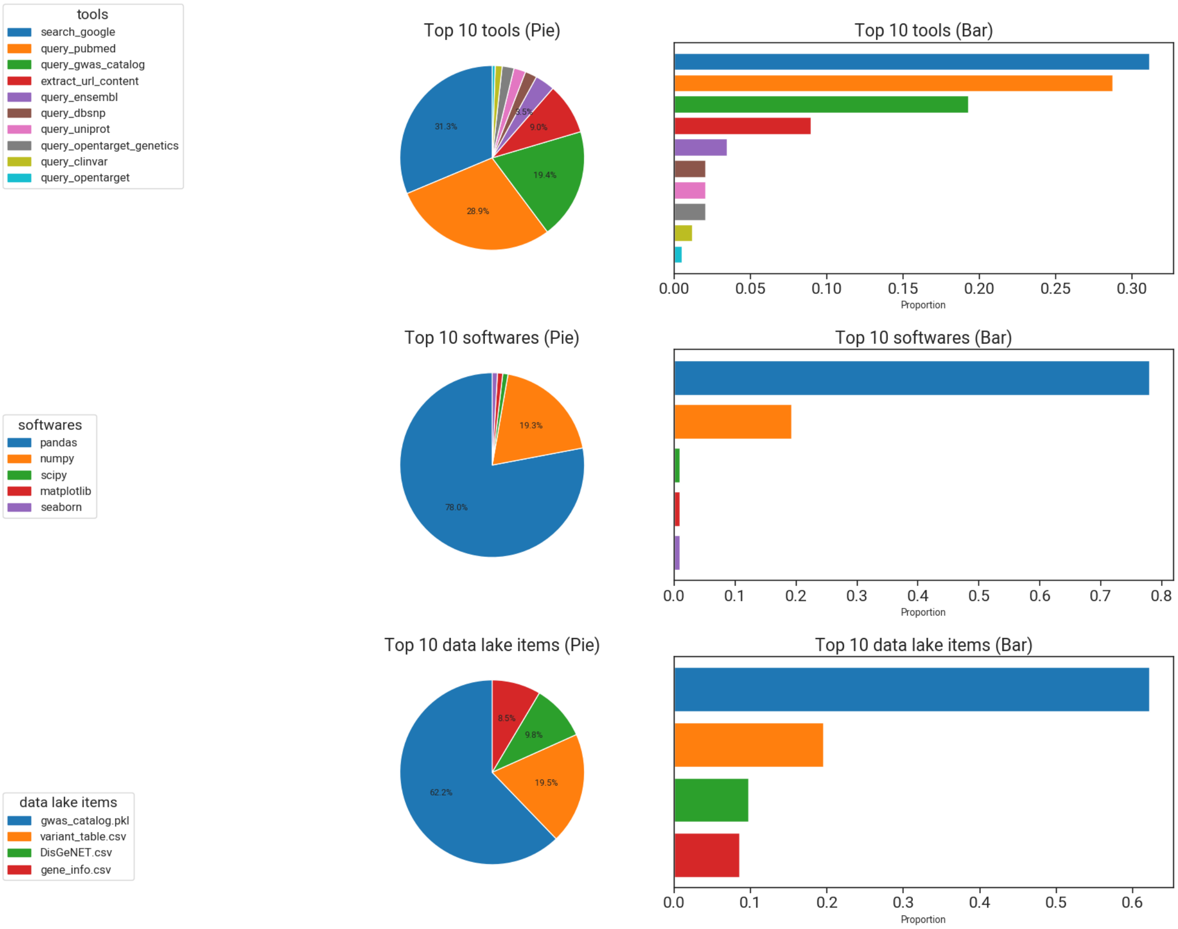


**Figure 16:** Detailed analysis on the tools, datasets, and software used for variant prioritization task.

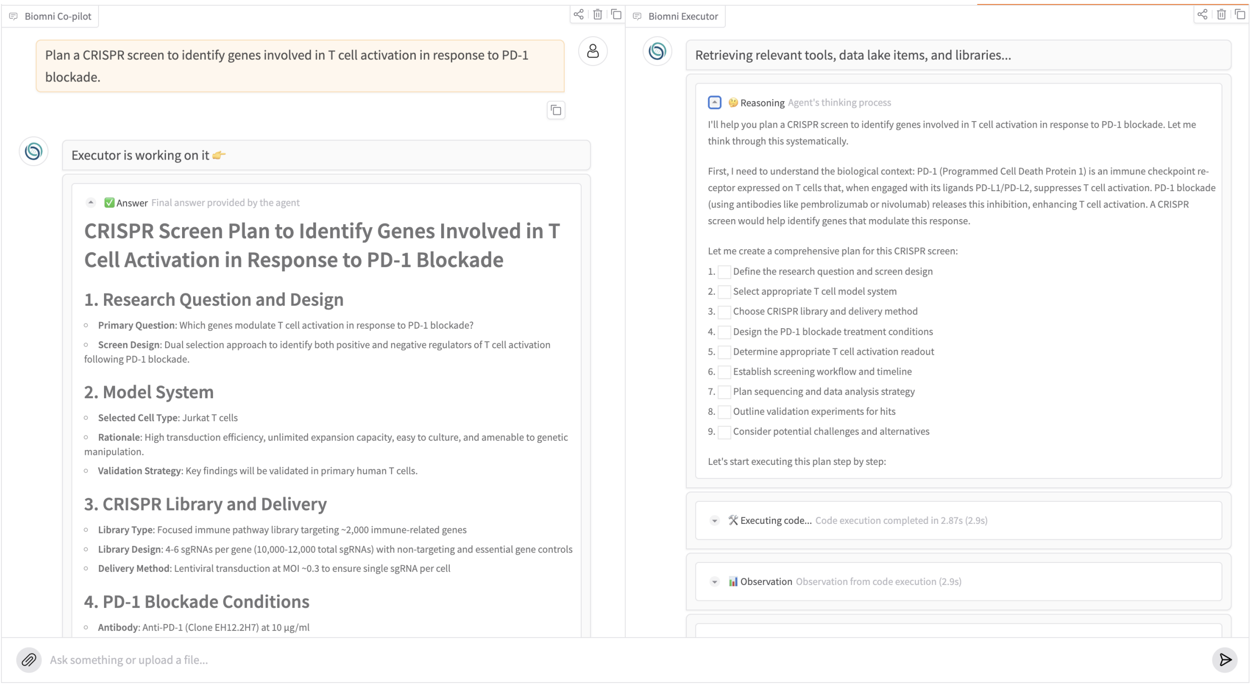


**Figure 17:** Biomni user interface.

#### Supplementary Tables

**Table 1:** Tool descriptions for `biomni.literature`

| **Tool Name** | **Tool Description** |
|---|---|
| `fetch_supplementary_info_from_doi` | Fetches supplementary information for a paper given its DOI and returns a research log. |
| `query_arxiv` | Query arXiv for papers based on the provided search query and return formatted search results. |
| `query_scholar` | Query Google Scholar for papers based on the provided search query and return the first search result. |
| `query_pubmed` | Query PubMed for papers based on the provided search query and return formatted results. |
| `search_google` | Search the web using Google search and return results with title, URL, and description. |
| `extract_url_content` | Extract the text content of a webpage using requests and BeautifulSoup, removing unwanted elements and formatting the result. |
| `extract_pdf_content` | Extract text content from a PDF file given its URL. |

**Table 2:** Tool descriptions for biomni.biochemistry

| Tool Name | Tool Description |
|---|---|
| analyze_circular_dichroism_spectra | Analyzes circular dichroism (CD) spectroscopy data to determine secondary structure and thermal stability of biomolecules. |
| analyze_rna_secondary_structure_features | Calculate numeric values for various structural features of an RNA secondary structure in dot-bracket notation. |
| analyze_protease_kinetics | Analyze protease kinetics data from fluorogenic peptide cleavage assays, fits the data to Michaelis-Menten kinetics, and determines key kinetic parameters (kcat, KM, and catalytic efficiency). |
| analyze_enzyme_kinetics_assay | Performs in vitro enzyme kinetics assay and analyzes the dose-dependent effects of modulators. |
| analyze_itc_binding_thermodynamics | Analyzes isothermal titration calorimetry (ITC) data to determine binding affinity and thermodynamic parameters. |
| analyze_protein_conservation | Perform multiple sequence alignment and phylogenetic analysis to identify conserved protein regions. |

**Table 3:** Tool descriptions for biomni.bioengineering

| Tool Name | Tool Description |
|---|---|
| analyze_cell_migration_metrics | Analyze cell migration metrics from time-lapse microscopy images and generate quantitative measurements of cell movement. |
| perform_crispr_cas9_genome_editing | Simulates CRISPR-Cas9 genome editing process including guide RNA design, delivery, and analysis. |
| analyze_calcium_imaging_data | Analyze calcium imaging data to quantify neuronal activity metrics including cell counts, event rates, decay times, and signal-to-noise ratios. |
| analyze_in_vitro_drug_release_kinetics | Analyzes in vitro drug release kinetics from biomaterial formulations and determines the best fitting kinetic model. |
| analyze_myofiber_morphology | Quantifies morphological properties of myofibers in microscopy images of tissue sections. |
| decode_behavior_from_neural_trajectories | Model neural activity trajectories and decode behavioral variables from neural spiking data. |
| simulate_whole_cell_ode_model | Simulate a whole-cell model represented as a system of ordinary differential equations (ODEs). |

**Table 4:** Tool descriptions for `biomni.biophysics`

| Tool Name | Tool Description |
|---|---|
| `predict_protein_disorder_regions` | Predicts intrinsically disordered regions (IDRs) in a protein sequence using IUPred2A algorithm. |
| `analyze_cell_morphology_and_cytoskeleton` | Quantifies cell morphology and cytoskeletal organization from fluorescence microscopy images. |
| `analyze_tissue_deformation_flow` | Quantify tissue deformation and flow dynamics from microscopy image sequence. |

**Table 5:** Tool descriptions for `biomni.cancer_biology`

| Tool Name | Tool Description |
|---|---|
| `analyze_ddr_network_in_cancer` | Analyze DNA Damage Response (DDR) network alterations and dependencies in cancer samples by reconstructing the DDR network from genomic data, identifying disruptions, and analyzing dependencies between DDR pathway components. |
| `analyze_cell_senescence_and_apoptosis` | Analyze flow cytometry data to quantify senescent and apoptotic cell populations from FCS files. |
| `detect_and_annotate_somatic_mutations` | Detects and annotates somatic mutations in tumor samples compared to matched normal samples using GATK Mutect2 for variant calling, GATK FilterMutectCalls for filtering, and SnpEff for functional annotation. |
| `detect_and_characterize_structural_variations` | Detects and characterizes structural variations (SVs) in genomic sequencing data using LUMPY for SV detection followed by annotation with COSMIC and/or ClinVar databases. |
| `perform_gene_expression_nmf_analysis` | Performs Non-negative Matrix Factorization (NMF) on gene expression data to extract metagenes and their associated sample weights for tumor subtype identification. |

**Table 6:** Tool descriptions for `biomni.cell_biology`

| Tool Name | Tool Description |
|---|---|
| `quantify_cell_cycle_phases_from_microscopy` | Quantify the percentage of cells in each cell cycle phase (G1, S, G2/M) using Calcofluor white stained microscopy images by segmenting cells and analyzing their morphological features. |
| `quantify_and_cluster_cell_motility` | Quantify cell motility features from time-lapse microscopy images and cluster cells based on motility patterns. |
| `perform_facs_cell_sorting` | Performs Fluorescence-Activated Cell Sorting (FACS) to enrich cell populations based on fluorescence characteristics. |
| `analyze_flow_cytometry_immunophenotyping` | Analyze flow cytometry data to identify and quantify specific cell populations based on surface markers. |
| `analyze_mitochondrial_morphology_and_potential` | Quantifies metrics of mitochondrial morphology and membrane potential from fluorescence microscopy images. |

**Table 7:** Tool descriptions for biomni.molecular_biology

| Tool Name | Tool Description |
|---|---|
| annotate_open_reading_frames | Find all Open Reading Frames (ORFs) in a DNA sequence using Biopython. Searches both forward and reverse complement strands if specified. |
| annotate_plasmid | Annotate a DNA sequence using pLannotate's command-line interface to identify features such as genes, promoters, and origins of replication. |
| get_gene_coding_sequence | Retrieves the coding sequence(s) of a specified gene from NCBI Entrez. |
| get_plasmid_sequence | Retrieves plasmid sequences from either Addgene or NCBI based on the provided identifier. |
| align_sequences | Align short sequences (primers) to a longer sequence, allowing for one mismatch. Checks both forward and reverse complement strands. |
| pcr_simple | Simulate PCR amplification with given primers and sequence, returning products and binding details. |
| pcr_complex_multi_primers | Simulate PCR amplification with multiple primers, considering all possible primer combinations and their potential products. |
| digest_sequence | Simulates restriction enzyme digestion of a DNA sequence and returns the resulting fragments with their properties. |
| golden_gate | Simulate a GoldenGate cloning reaction with Type IIS restriction enzymes to predict assembly products. |
| oligo_assembly | Assemble two DNA sequences into an oligo with overhangs. Automatically detects overhang type and length. |
| gibson_assembly | Simulate a Gibson Assembly reaction to join DNA fragments with overlapping regions. |
| find_restriction_sites | Identifies restriction enzyme sites in a given DNA sequence for specified enzymes. |
| find_restriction_enzymes | Finds common restriction enzyme sites in a DNA sequence. |
| design_primers_with_overhangs | Design two primers to amplify a target sequence with optional overhangs. |
| find_sequence_mutations | Compare query sequence against reference sequence to identify mutations. |
| get_molecular_cloning_instructions | Returns a dictionary containing molecular cloning instructions and important notes. |
| calculate_element_distances | Calculate pairwise distances between elements on a DNA sequence/plasmid, providing forward, reverse (for circular sequences), and shortest path distances. |

**Table 8:** Continue tool descriptions for biomni.molecular_biology

| Tool Name | Tool Description |
|---|---|
| design_knockout_sgrna | Design sgRNAs for CRISPR knockout by searching precomputed sgRNA libraries for a specific gene. |
| design_golden_gate_oligos | Design complementary oligonucleotides with Type IIS restriction enzyme overhangs for Golden Gate assembly based on restriction site analysis of the backbone. |
| get_oligo_annealing_protocol | Return a standard protocol for annealing complementary oligonucleotides without phosphorylation. |
| get_golden_gate_assembly_protocol | Return a customized protocol for Golden Gate assembly based on the number of inserts and specific DNA sequences. |
| get_bacterial_transformation_protocol | Return a standard protocol for bacterial transformation with detailed steps. |
| design_primer | Design a single primer within a given DNA sequence window based on GC content and melting temperature constraints. |
| design_verification_primers | Design Sanger sequencing primers to verify a specific region in a plasmid, using existing primers when possible and designing new ones as needed. |

**Table 9:** Tool descriptions for biomni.genetics

| Tool Name | Tool Description |
|---|---|
| liftover_coordinates | Perform liftover of genomic coordinates between hg19 and hg38 genome builds with detailed step-by-step explanations. |
| bayesian_finemapping_with_deep_vi | Performs Bayesian fine-mapping from GWAS summary statistics using deep variational inference to compute posterior inclusion probabilities and credible sets for putative causal variants. |
| analyze_cas9_mutation_outcomes | Analyzes and categorizes mutations induced by Cas9 at target sites, generating detailed statistics on mutation types. |
| analyze_crispr_genome_editing | Analyzes CRISPR-Cas9 genome editing results by comparing original and edited sequences to identify mutations and characterize edited loci. |
| simulate_demographic_history | Simulate DNA sequences with specified demographic and coalescent histories using msprime. |
| identify_transcription_factor_binding_sites | Identifies binding sites for a specific transcription factor in a genomic sequence using position weight matrices from the JASPAR database. |
| fit_genomic_prediction_model | Fit a linear mixed model for genomic prediction using genotype and phenotype data. |
| perform_pcr_and_gel_electrophoresis | Performs PCR amplification of a target transgene and visualizes results using agarose gel electrophoresis. |
| analyze_protein_phylogeny | Perform phylogenetic analysis on a set of protein sequences, including multiple sequence alignment, tree construction, and visualization. |

**Table 10:** Tool descriptions for `biomni.genomics`

| Tool Name | Tool Description |
|---|---|
| annotate_celltype_scRNA | Annotate cell types in single-cell RNA-seq data based on gene markers and transferred labels using LLM. |
| create_scvi_embeddings_scRNA | Creates scVI and scANVI embeddings for single-cell RNA-seq data, training models and saving the results to a new AnnData object. |
| create_harmony_embeddings_scRNA | Performs batch effect correction on single-cell RNA-seq data using the Harmony algorithm and saves the integrated embeddings. |
| get_uce_embeddings_scRNA | Generate UCE (Universal Cell Embeddings) for single-cell RNA sequencing data to enable cell type identification and mapping to reference datasets. |
| map_to_ima_interpret_scRNA | Map cell embeddings from the input dataset to the Integrated Megascale Atlas reference dataset using UCE embeddings for cell type annotation. |
| get_rna_seq_archs4 | Given a gene name, fetch and return RNA-seq expression data (transcripts-per-million) across tissues from the ARCHS4 database. |
| get_gene_set_enrichment_analysis_supported_database_list | Returns a list of supported databases available for gene set enrichment analysis |
| gene_set_enrichment_analysis | Perform enrichment analysis for a list of genes to identify pathways, transcription factors, or other biological relationships. |
| analyze_chromatin_interactions | Analyze chromatin interactions from Hi-C data to identify enhancer-promoter interactions and topologically associated domains (TADs). |
| analyze_comparative_genomics_and_haplotypes | Perform comparative genomics and haplotype analysis on multiple genome samples. Aligns genomes to a reference, identifies variants, analyzes shared and unique genomic regions, and determines haplotype structure. |
| perform_chipseq_peak_calling_with_macs2 | Perform ChIP-seq peak calling using MACS2 to identify genomic regions with significant binding. |
| find_enriched_motifs_with_homer | Find DNA sequence motifs enriched in genomic regions using the HOMER motif discovery software. |
| analyze_genomic_region_overlap | Analyze overlaps between two or more sets of genomic regions and generate a research log summarizing the analysis. |

**Table 11:** Tool descriptions for `biomni.immunology`

| **Tool Name** | **Tool Description** |
|---|---|
| `analyze_atac_seq_differential_accessibility` | Perform ATAC-seq peak calling and differential accessibility analysis using MACS2. |
| `analyze_bacterial_growth_curve` | Analyzes bacterial growth curve data to determine growth parameters like doubling time, growth rate, and lag phase. |
| `isolate_purify_immune_cells` | Isolates and purifies immune cells from tissue samples and returns a research log of the process. |
| `estimate_cell_cycle_phase_durations` | Estimate cell cycle phase durations using dual-nucleoside pulse labeling data and mathematical modeling. |
| `track_immune_cells_under_flow` | Track immune cells under flow conditions and classify their behaviors. |
| `analyze_cfse_cell_proliferation` | Analyze CFSE-labeled cell samples to quantify cell division and proliferation from flow cytometry data. |
| `analyze_cytokine_production_in_cd4_tcells` | Analyze cytokine production (IFN-$\gamma$, IL-17) in CD4+ T cells after antigen stimulation using flow cytometry data. |
| `analyze_ebv_antibody_titers` | Analyze ELISA data to quantify EBV antibody titers in plasma/serum samples. |
| `analyze_cns_lesion_histology` | Analyzes histological images of CNS lesions to quantify immune cell infiltration, demyelination, and tissue damage. |
| `analyze_immunohistochemistry_image` | Analyzes immunohistochemistry images to quantify protein expression and spatial distribution. |

**Table 12:** Tool descriptions for `biomni.microbiology`

| **Tool Name** | **Tool Description** |
|---|---|
| `optimize_anaerobic_digestion_process` | Optimize anaerobic digestion process conditions to maximize VFA production or methane yield. |
| `analyze_arsenic_speciation_hplc_icpms` | Analyzes arsenic speciation in liquid samples using HPLC-ICP-MS technique and returns a detailed research log. |
| `count_bacterial_colonies` | Count bacterial colonies from an image of agar plate using computer vision techniques. |
| `annotate_bacterial_genome` | Annotate a bacterial genome using Prokka to identify genes, proteins, and functional features. |
| `enumerate_bacterial_cfu_by_serial_dilution` | Quantify bacterial concentration (CFU/mL) using serial dilutions and spot plating. |
| `model_bacterial_growth_dynamics` | Model bacterial population dynamics over time using ordinary differential equations. |
| `quantify_biofilm_biomass_crystal_violet` | Quantifies biofilm biomass using crystal violet staining assay data and generates a detailed research log of the analysis. |
| `segment_and_analyze_microbial_cells` | Perform automated cell segmentation and quantify morphological metrics from fluorescence microscopy images. |
| `segment_cells_with_deep_learning` | Perform cell segmentation on fluorescence microscopy images using deep learning models from the Cellpose/Omnipose library. |
| `simulate_generalized_lotka_volterra_dynamics` | Simulate microbial community dynamics using the Generalized Lotka-Volterra (gLV) model. |
| `predict_rna_secondary_structure` | Predict the secondary structure of an RNA molecule using ViennaRNA and generate visualization files. |
| `simulate_microbial_population_dynamics` | Performs stochastic simulation of microbial population dynamics using the Gillespie algorithm. |

**Table 13:** Tool descriptions for biomni.pathology

| Tool Name | Tool Description |
|---|---|
| analyze_aortic_diameter_and_geometry | Analyze aortic diameter and geometry from cardiovascular imaging data to measure aortic root diameter, ascending aorta diameter, and calculate geometric parameters such as tortuosity and dilation indices. |
| analyze_atp_luminescence_assay | Analyze luminescence-based ATP assay data to determine intracellular ATP concentration and generate a detailed research log of the analysis. |
| analyze_thrombus_histology | Analyze histological images of thrombus samples stained with H&E to identify and quantify different thrombus components (fresh, cellular lysis, endothelialization, fibroblastic reaction). |
| analyze_intracellular_calcium_with_rhod2 | Analyzes intracellular calcium concentration using Rhod-2 fluorescent indicator from microscopy images. |
| quantify_corneal_nerve_fibers | Quantify the volume/density of immunofluorescence-labeled corneal nerve fibers from microscopy images. |
| segment_and_quantify_cells_in_multiplexed_images | Segment cells and quantify protein expression levels from multichannel tissue images. |
| analyze_bone_microct_morphometry | Analyze bone microarchitecture parameters from 3D micro-CT images, calculating metrics such as bone mineral density (BMD), bone volume (BV), trabecular number (Tb.N), trabecular thickness (Tb.Th), and trabecular separation (Tb.S). |

**Table 14:** Tool descriptions for biomni.pharmacology

| Tool Name | Tool Description |
|---|---|
| run_diffdock_with_smiles | Run DiffDock molecular docking simulation using a protein structure and a SMILES string for the ligand. Uses Docker to execute the DiffDock algorithm. |
| docking_autodock_vina | Performs molecular docking using AutoDock Vina to predict binding affinities between small molecules and a receptor protein. |
| run_autosite | Runs AutoSite on a protein structure to identify potential binding sites and returns a research log with the results. |
| retrieve_topk_repurposing_drugs_from_disease_txgnn | Computes TxGNN model predictions for drug repurposing for a given disease and returns the top predicted drugs with their scores. |
| predict_admet_properties | Predicts ADMET (Absorption, Distribution, Metabolism, Excretion, Toxicity) properties for a list of compounds using pretrained models. |
| predict_binding_affinity_protein_1d_sequence | Predicts the binding affinity between small molecules and a protein sequence using pre-trained deep learning models. |
| analyze_accelerated_stability_of_pharmaceutical_formulations | Analyzes the stability of pharmaceutical formulations under accelerated storage conditions and generates a research log of the results. |
| run_3d_chondrogenic_aggregate_assay | Generates a detailed protocol for performing a 3D chondrogenic aggregate culture assay to evaluate compounds' effects on chondrogenesis. |
| grade_adverse_events_using_vcog_ctcae | Grade and monitor adverse events in animal studies using the VCOG-CTCAE standard. |
| analyze_radiolabeled_antibody_biodistribution | Analyze biodistribution and pharmacokinetic profile of radiolabeled antibodies, including tissue distribution, half-lives, and tumor-to-normal tissue ratios. |
| estimate_alpha_particle_radiotherapy_dosimetry | Estimate radiation absorbed doses to tumor and normal organs for alpha-particle radiotherapeutics using the Medical Internal Radiation Dose (MIRD) schema. |
| perform_mwas_cyp2c19_metabolizer_status | Perform a Methylome-wide Association Study (MWAS) to identify CpG sites significantly associated with CYP2C19 metabolizer status. |
| calculate_physicochemical_properties | Calculate key physicochemical properties of a drug candidate molecule including molecular weight, cLogP, TPSA, H-bond donors/acceptors, and other drug-like characteristics. |
| analyze_xenograft_tumor_growth_inhibition | Analyze tumor growth inhibition in xenograft models across different treatment groups. |
| analyze_western_blot | Performs densitometric analysis of Western blot images to quantify relative protein expression. |

**Table 15:** Tool descriptions for biomni.physiology

| Tool Name | Tool Description |
|---|---|
| reconstruct_3d_face_from_mri | Generate a 3D model of facial anatomy from MRI scans of the head and neck. |
| analyze_abr_waveform_p1_metrics | Extracts P1 amplitude and latency from Auditory Brainstem Response (ABR) waveform data. P1 (Wave I) is typically the first positive peak in the ABR waveform and is a critical marker for auditory function assessment. |
| analyze_ciliary_beat_frequency | Analyze ciliary beat frequency from high-speed video microscopy data using FFT analysis. |
| analyze_protein_colocalization | Analyze colocalization between two fluorescently labeled proteins in microscopy images. |
| perform_cosinor_analysis | Performs cosinor analysis on physiological time series data to characterize circadian rhythms. |
| calculate_brain_adc_map | Calculate Apparent Diffusion Coefficient (ADC) map from diffusion-weighted MRI data using the monoexponential diffusion model. |
| analyze_endolysosomal_calcium_dynamics | Analyze calcium dynamics in endo-lysosomal compartments using ELGA/ELGA1 probe data. |
| analyze_fatty_acid_composition_by_gc | Analyzes fatty acid composition in tissue samples using gas chromatography data. |
| analyze_hemodynamic_data | Analyzes raw blood pressure data to calculate key hemodynamic parameters including systolic blood pressure, diastolic blood pressure, mean arterial pressure, and heart rate. |
| simulate_thyroid_hormone_pharmacokinetics | Simulates the transport and binding of thyroid hormones across different tissue compartments using an ODE-based pharmacokinetic model. |
| quantify_amyloid_beta_plaques | Analyzes an image to detect, quantify, and characterize amyloid-beta plaques commonly found in Alzheimer's disease tissue samples. |

**Table 16:** Tool descriptions for biomni.synthetic_biology

| Tool Name | Tool Description |
|---|---|
| engineer_bacterial_genome_for_therapeutic_delivery | Engineer a bacterial genome by integrating therapeutic genetic parts for therapeutic delivery. |
| analyze_bacterial_growth_rate | Analyze bacterial growth data and extract growth parameters from OD600 measurements. |
| analyze_barcode_sequencing_data | Analyze sequencing data to extract, quantify and determine lineage relationships of barcodes. |
| analyze_bifurcation_diagram | Performs bifurcation analysis on a dynamical system and generates a bifurcation diagram. |
| create_biochemical_network_sbml_model | Generate a mathematical model of a biochemical network in SBML format. |
| optimize_codons_for_heterologous_expression | Analyzes and optimizes a DNA/RNA sequence for improved expression in a heterologous host organism. |
| simulate_gene_circuit_with_growth_feedback | Simulate gene regulatory circuit dynamics with growth feedback, tracking gene expression levels and cell growth over time. |
| identify_fas_functional_domains | Identifies functional domains within a Fatty Acid Synthase (FAS) sequence and predicts their roles. |

**Table 17:** Tool descriptions for biomni.systems_biology

| Tool Name | Tool Description |
|---|---|
| perform_flux_balance_analysis | Perform Flux Balance Analysis (FBA) on a genome-scale metabolic network model to predict metabolic flux distributions. |
| model_protein_dimerization_network | Model protein dimerization networks to find equilibrium concentrations of dimers based on monomer concentrations and binding affinities. |
| simulate_metabolic_network_perturbation | Construct and simulate kinetic models of metabolic networks and analyze their responses to perturbations. |
| simulate_protein_signaling_network | Simulate protein signaling network dynamics using ODE-based logic modeling with normalized Hill functions. |
| compare_protein_structures | Compares two protein structures to identify structural differences and conformational changes. |
| simulate_renin_angiotensin_system_dynamics | Simulate the time-dependent concentrations of renin-angiotensin system (RAS) components. |

**Table 18:** Tool descriptions for biomni.support_tools

| Tool Name | Tool Description |
|---|---|
| run_python_repl | Executes a Python command in the notebook environment and returns the output as a string. |
| read_function_source_code | Read the source code of a function from any module path. |

**Table 19:** Tool descriptions for `biomni.database`

| **Tool Name** | **Tool Description** |
|---|---|
| `query_uniprot` | Query the UniProt REST API using either natural language or a direct endpoint to retrieve protein information. |
| `query_alphafold` | Query the AlphaFold Database API for protein structure predictions and information. |
| `query_interpro` | Query the InterPro REST API using natural language or a direct endpoint to retrieve information about protein domains or families. |
| `query_pdb` | Query the RCSB PDB database using natural language or a direct structured query to find protein structures. |
| `query_pdb_identifiers` | Retrieve detailed data and/or download files for PDB identifiers. |
| `query_kegg` | Take a natural language prompt and convert it to a structured KEGG API query, then execute the query. |
| `query_stringdb` | Query the STRING protein interaction database using natural language or direct endpoint. |
| `query_iucn` | Query the IUCN Red List API using natural language or a direct endpoint to retrieve species conservation status information. |
| `query_paleobiology` | Query the Paleobiology Database (PBDB) API using natural language or a direct endpoint. |
| `query_jaspar` | Query the JASPAR REST API using natural language or a direct endpoint to retrieve transcription factor binding profiles. |
| `query_worms` | Query the World Register of Marine Species (WoRMS) REST API using natural language or a direct endpoint. |
| `query_cbioportal` | Query the cBioPortal REST API using natural language or a direct endpoint to access cancer genomics data. |
| `query_clinvar` | Take a natural language prompt and convert it to a structured ClinVar query to search for genetic variants. |
| `query_geo` | Query the NCBI Gene Expression Omnibus (GEO) using natural language or a direct search term. |
| `query_dbsnp` | Query the NCBI dbSNP database using natural language or a direct search term. |
| `query_ucsc` | Query the UCSC Genome Browser API using natural language or a direct endpoint. |

**Table 20:** Continue tool descriptions for `biomni.database`

| Tool Name | Tool Description |
|---|---|
| query_ensembl | Query the Ensembl REST API using natural language or a direct endpoint to retrieve genomic data. |
| query_opentarget_genetics | Query the OpenTargets Genetics API using natural language or a direct GraphQL query to retrieve information about genetic targets and variants. |
| query_opentarget | Query the OpenTargets Platform API using natural language or a direct GraphQL query to access drug targets, diseases, and mechanisms data. |
| query_gwas_catalog | Query the GWAS Catalog API using natural language or a direct endpoint to retrieve genetic association studies data. |
| query_gnomad | Query gnomAD for variants in a gene using natural language or direct gene symbol. |
| blast_sequence | Identifies a DNA or protein sequence using NCBI BLAST and returns information about the best alignment. |
| query_reactome | Query the Reactome database using natural language or a direct endpoint to retrieve information about biological pathways. |
| query_regulomedb | Query the RegulomeDB database using natural language or direct endpoint specification to get information about regulatory elements. |
| query_pride | Query the PRIDE (PRoteomics IDEntifications) database using natural language or a direct endpoint. |
| query_gtopdb | Query the Guide to PHARMACOLOGY database (GtoPdb) using natural language or a direct endpoint. |
| region_to_ccre_screen | Retrieves information about candidate cis-regulatory elements (cCREs) that intersect with a specified genomic region. |
| get_genes_near_ccre | Identifies the nearest genes to a specified candidate cis-Regulatory Element (cCRE) by querying the SCREEN database. |
| query_remap | Query the ReMap database for regulatory elements and transcription factor binding sites using natural language or direct API endpoints. |
| query_mpd | Query the Mouse Phenome Database (MPD) for mouse strain phenotype data using natural language or direct endpoint access. |
| query_emdb | Query the Electron Microscopy Data Bank (EMDB) for 3D macromolecular structures using natural language or direct endpoint access. |

**Table 21:** Data lake descriptions

| File Name | Description |
|---|---|
| affinity_capture-ms.csv | Protein-protein interactions detected via affinity capture and mass spectrometry. |
| affinity_capture-rna.csv | Protein-RNA interactions detected by affinity capture. |
| BindingDB_All_202409.tsv | Measured binding affinities between proteins and small molecules for drug discovery. |
| broad_repurposing_hub_molecule_with_smiles.csv | Molecules from Broad Institute's Drug Repurposing Hub with SMILES annotations. |
| broad_repurposing_hub_phase_moa_target_info.csv | Drug phases, mechanisms of action, and target information from Broad Institute. |
| co-fractionation.csv | Protein-protein interactions from co-fractionation experiments. |
| Cosmic_Breakpoints_v101_GRCh38.csv | Genomic breakpoints associated with cancers from COSMIC database. |
| Cosmic_CancerGeneCensusHallmarksOfCancer_v101_GRCh38.csv | Hallmarks of cancer genes from COSMIC. |
| Cosmic_CancerGeneCensus_v101_GRCh38.csv | Census of cancer-related genes from COSMIC. |
| Cosmic_ClassificationPaper_v101_GRCh38.csv | Cancer classifications and annotations from COSMIC. |
| Cosmic_Classification_v101_GRCh38.csv | Classification of cancer types from COSMIC. |
| Cosmic_CompleteCNA_v101_GRCh38.tsv.gz | Complete copy number alterations data from COSMIC. |
| Cosmic_CompleteDifferentialMethylation_v101_GRCh38.tsv.gz | Differential methylation patterns from COSMIC. |
| Cosmic_CompleteGeneExpression_v101_GRCh38.tsv.gz | Gene expression data across cancers from COSMIC. |
| Cosmic_Fusion_v101_GRCh38.csv | Gene fusion events from COSMIC. |
| Cosmic_Genes_v101_GRCh38.csv | List of genes associated with cancer from COSMIC. |
| Cosmic_GenomeScreensMutant_v101_GRCh38.tsv.gz | Genome screening mutations from COSMIC. |
| Cosmic_MutantCensus_v101_GRCh38.csv | Catalog of cancer-related mutations from COSMIC. |
| Cosmic_ResistanceMutations_v101_GRCh38.csv | Resistance mutations related to therapeutic interventions from COSMIC. |
| czi_census_datasets_v4.csv | Datasets from the Chan Zuckerberg Initiative's Cell Census. |
| DisGeNET.csv | Gene-disease associations from multiple sources. |
| dosage_growth_defect.csv | Gene dosage changes affecting growth. |
| enamine_cloud_library_smiles.pkl | Compounds from Enamine REAL library with SMILES annotations. |
| genebass_missense_LC_filtered.pkl | Filtered missense variants from GeneBass. |
| genebass_pLoF_filtered.pkl | Predicted loss-of-function variants from GeneBass. |
| genebass_synonymous_filtered.pkl | Filtered synonymous variants from GeneBass. |
| gene_info.csv | Comprehensive gene information. |
| genetic_interaction.csv | Genetic interactions between genes. |
| go-plus.json | Gene ontology data for functional gene annotations. |
| gtex_tissue_gene_tpm.csv | Gene expression (TPM) across human tissues from GTEx. |
| gwas_catalog.pkl | Genome-wide association studies (GWAS) results. |
| marker_celltype.csv | Cell type marker genes for identification. |
| McPAS-TCR.csv | T-cell receptor sequences and specificity data from McPAS database. |
| miRDB_v6.0_results.csv | Predicted microRNA targets from miRDB. |
| miRTarBase_microRNA_target_interaction.csv | Experimentally validated microRNA-target interactions from miRTarBase. |
| miRTarBase_microRNA_target_interaction_pubmed_abtract.txt | PubMed abstracts for microRNA-target interactions in miRTarBase. |
| miRTarBase_MicroRNA_Target_Sites.csv | Binding sites of microRNAs on target genes from miRTarBase. |

**Table 22:** Continue data lake descriptions

| File Name | Description |
|---|---|
| mousemine_m1_positional_geneset.csv | Positional gene sets from MouseMine. |
| mousemine_m2_curated_geneset.csv | Curated gene sets from MouseMine. |
| mousemine_m3_regulatory_target_geneset.csv | Regulatory target gene sets from MouseMine. |
| mousemine_m5_ontology_geneset.csv | Ontology-based gene sets from MouseMine. |
| mousemine_m8_celltype_signature_geneset.csv | Cell type signature gene sets from MouseMine. |
| mousemine_mh_hallmark_geneset.csv | Hallmark gene sets from MouseMine. |
| msigdb_human_c1_positional_geneset.csv | Human positional gene sets from MSigDB. |
| msigdb_human_c2_curated_geneset.csv | Curated human gene sets from MSigDB. |
| msigdb_human_c3_regulatory_target_geneset.csv | Regulatory target gene sets from MSigDB. |
| msigdb_human_c3_subset_transcription_factor_targets_from_GTRD.csv | Transcription factor targets from GTRD/MSigDB. |
| msigdb_human_c4_computational_geneset.csv | Computationally derived gene sets from MSigDB. |
| msigdb_human_c5_ontology_geneset.csv | Ontology-based gene sets from MSigDB. |
| msigdb_human_c6_oncogenic_signature_geneset.csv | Oncogenic signatures from MSigDB. |
| msigdb_human_c7_immunologic_signature_geneset.csv | Immunologic signatures from MSigDB. |
| msigdb_human_c8_celltype_signature_geneset.csv | Cell type signatures from MSigDB. |
| msigdb_human_h_hallmark_geneset.csv | Hallmark gene sets from MSigDB. |
| omim.csv | Genetic disorders and associated genes from OMIM. |
| proteinatlas.tsv | Protein expression data from Human Protein Atlas. |
| proximity_label-ms.csv | Protein interactions via proximity labeling and mass spectrometry. |
| reconstituted_complex.csv | Protein complexes reconstituted in vitro. |
| synthetic_growth_defect.csv | Synthetic growth defects from genetic interactions. |
| synthetic_lethality.csv | Synthetic lethal interactions. |
| synthetic_rescue.csv | Genetic interactions rescuing phenotypes. |
| two-hybrid.csv | Protein-protein interactions detected by yeast two-hybrid assays. |
| variant_table.csv | Annotated genetic variants table. |
| Virus-Host_PPI_P-HIPSTER_2020.csv | Virus-host protein-protein interactions from P-HIPSTER. |

**Table 23:** Software descriptions

| Software Name | Description |
|---|---|
| biopython | [Python Package] A set of tools for biological computation including parsers for bioinformatics files, access to online services, and interfaces to common bioinformatics programs. |
| biom-format | [Python Package] The Biological Observation Matrix (BIOM) format is designed for representing biological sample by observation contingency tables with associated metadata. |
| scanpy | [Python Package] A scalable toolkit for analyzing single-cell gene expression data, specifically designed for large datasets using AnnData. |
| scikit-bio | [Python Package] Data structures, algorithms, and educational resources for bioinformatics, including sequence analysis, phylogenetics, and ordination methods. |
| anndata | [Python Package] A Python package for handling annotated data matrices in memory and on disk, primarily used for single-cell genomics data. |
| mudata | [Python Package] A Python package for multimodal data storage and manipulation, extending AnnData to handle multiple modalities. |
| pyliftover | [Python Package] A Python implementation of UCSC liftOver tool for converting genomic coordinates between genome assemblies. |
| biopandas | [Python Package] A package that provides pandas DataFrames for working with molecular structures and biological data. |
| biotite | [Python Package] A comprehensive library for computational molecular biology, providing tools for sequence analysis, structure analysis, and more. |
| gget | [Python Package] A toolkit for accessing genomic databases and retrieving sequences, annotations, and other genomic data. |
| lifelines | [Python Package] A complete survival analysis library for fitting models, plotting, and statistical tests. |
| scvi-tools | [Python Package] A package for probabilistic modeling of single-cell omics data, including deep generative models. |

**Table 24:** Continue software descriptions

| Software Name | Description |
|---|---|
| gseapy | [Python Package] A Python wrapper for Gene Set Enrichment Analysis (GSEA) and visualization. |
| scrublet | [Python Package] A tool for detecting doublets in single-cell RNA-seq data. |
| cellxgene-census | [Python Package] A tool for accessing and analyzing the CellxGene Census, a collection of single-cell datasets. |
| hyperopt | [Python Package] A Python library for optimizing hyperparameters of machine learning algorithms. |
| scvelo | [Python Package] A tool for RNA velocity analysis in single cells using dynamical models. |
| pysam | [Python Package] A Python module for reading, manipulating and writing genomic data sets in SAM/BAM/VCF/BCF formats. |
| pyfaidx | [Python Package] A Python package for efficient random access to FASTA files. |
| pyranges | [Python Package] A Python package for interval manipulation with a pandas-like interface. |
| pybedtools | [Python Package] A Python wrapper for Aaron Quinlan's BEDTools programs. |
| rdkit | [Python Package] A collection of cheminformatics and machine learning tools for working with chemical structures and drug discovery. |
| deeppurpose | [Python Package] A deep learning library for drug-target interaction prediction and virtual screening. |
| pyscreener | [Python Package] A Python package for virtual screening of chemical compounds. |
| openbabel | [Python Package] A chemical toolbox designed to speak the many languages of chemical data, supporting file format conversion and molecular modeling. |
| descriptastorus | [Python Package] A library for computing molecular descriptors for machine learning applications in drug discovery. |

**Table 25:** Continue software descriptions

| Software Name | Description |
|---|---|
| `pymol` | [Python Package] A molecular visualization system for rendering and animating 3D molecular structures. |
| `openmm` | [Python Package] A toolkit for molecular simulation using high-performance GPU computing. |
| `pytdc` | [Python Package] A Python package for Therapeutics Data Commons, providing access to machine learning datasets for drug discovery. |
| `pandas` | [Python Package] A fast, powerful, and flexible data analysis and manipulation library for Python. |
| `numpy` | [Python Package] The fundamental package for scientific computing with Python, providing support for arrays, matrices, and mathematical functions. |
| `scipy` | [Python Package] A Python library for scientific and technical computing, including modules for optimization, linear algebra, integration, and statistics. |
| `scikit-learn` | [Python Package] A machine learning library featuring various classification, regression, and clustering algorithms. |
| `matplotlib` | [Python Package] A comprehensive library for creating static, animated, and interactive visualizations in Python. |
| `seaborn` | [Python Package] A statistical data visualization library based on matplotlib with a high-level interface for drawing attractive statistical graphics. |
| `statsmodels` | [Python Package] A Python module for statistical modeling and econometrics, including descriptive statistics and estimation of statistical models. |
| `pymc3` | [Python Package] A Python package for Bayesian statistical modeling and probabilistic machine learning. |
| `pystan` | [Python Package] A Python interface to Stan, a platform for statistical modeling and high-performance statistical computation. |
| `umap-learn` | [Python Package] Uniform Manifold Approximation and Projection, a dimension reduction technique. |

**Table 26:** Continue software descriptions

| **Software Name** | **Description** |
|---|---|
| faiss-cpu | [Python Package] A library for efficient similarity search and clustering of dense vectors. |
| harmony-pytorch | [Python Package] A PyTorch implementation of the Harmony algorithm for integrating single-cell data. |
| tiledb | [Python Package] A powerful engine for storing and analyzing large-scale genomic data. |
| tiledbsoma | [Python Package] A library for working with the SOMA (Stack of Matrices) format using TileDB. |
| h5py | [Python Package] A Python interface to the HDF5 binary data format, allowing storage of large amounts of numerical data. |
| tqdm | [Python Package] A fast, extensible progress bar for loops and CLI applications. |
| joblib | [Python Package] A set of tools to provide lightweight pipelining in Python, including transparent disk-caching and parallel computing. |
| opencv-python | [Python Package] OpenCV library for computer vision tasks, useful for image analysis in biological contexts. |
| PyPDF2 | [Python Package] A library for working with PDF files, useful for extracting text from scientific papers. |
| googlesearch-python | [Python Package] A library for performing Google searches programmatically. |
| scikit-image | [Python Package] A collection of algorithms for image processing in Python. |
| pymed | [Python Package] A Python library for accessing PubMed articles. |
| arxiv | [Python Package] A Python wrapper for the arXiv API, allowing access to scientific papers. |

**Table 27:** Continue software descriptions

| Software Name | Description |
|---|---|
| scholarly | [Python Package] A module to retrieve author and publication information from Google Scholar. |
| cryosparc-tools | [Python Package] Tools for working with cryoSPARC, a platform for cryo-EM data processing. |
| mageck | [Python Package] Analysis of CRISPR screen data. |
| igraph | [Python Package] Network analysis and visualization. |
| pyscenic | [Python Package] Analysis of single-cell RNA-seq data and gene regulatory networks. |
| cooler | [Python Package] Storage and analysis of Hi-C data. |
| trackpy | [Python Package] Particle tracking in images and video. |
| flowcytometrytools | [Python Package] Analysis and visualization of flow cytometry data. |
| cellpose | [Python Package] Cell segmentation in microscopy images. |
| viennarna | [Python Package] RNA secondary structure prediction. |
| PyMassSpec | [Python Package] Mass spectrometry data analysis. |
| python-libsbml | [Python Package] Working with SBML files for computational biology. |
| cobra | [Python Package] Constraint-based modeling of metabolic networks. |
| reportlab | [Python Package] Creation of PDF documents. |
| flowkit | [Python Package] Toolkit for processing flow cytometry data. |
| hmmlearn | [Python Package] Hidden Markov model analysis. |
| msprime | [Python Package] Simulation of genetic variation. |
| tskit | [Python Package] Handling tree sequences and population genetics data. |
| cyvcf2 | [Python Package] Fast parsing of VCF files. |
| pykalman | [Python Package] Kalman filter and smoother implementation. |

Table 28: Continue software descriptions

| Software Name | Description |
|---|---|
| fanc | [Python Package] Analysis of chromatin conformation data. |
| ggplot2 | [R Package] A system for declaratively creating graphics, based on The Grammar of Graphics. Use with subprocess.run(['Rscript', '-e', 'library(ggplot2); ...']). |
| dplyr | [R Package] A grammar of data manipulation, providing a consistent set of verbs that help you solve the most common data manipulation challenges. Use with subprocess. |
| tidyr | [R Package] A package that helps you create tidy data, where each column is a variable, each row is an observation, and each cell is a single value. Use with subprocess. |
| readr | [R Package] A fast and friendly way to read rectangular data like CSV, TSV, and FWF. Use with subprocess.run(['Rscript', '-e', 'library(readr); ...']). |
| stringr | [R Package] A cohesive set of functions designed to make working with strings as easy as possible. Use with subprocess calls. |
| Matrix | [R Package] A package that provides classes and methods for dense and sparse matrices. Required for Seurat. Use with subprocess calls. |
| Rcpp | [R Package] Seamless R and C++ Integration, allowing R functions to call compiled C++ code. Use with subprocess calls. |
| devtools | [R Package] Tools to make developing R packages easier, including functions to install packages from GitHub. Use with subprocess calls. |
| remotes | [R Package] Install R packages from GitHub, GitLab, Bitbucket, or other remote repositories. Use with subprocess calls. |
| DESeq2 | [R Package] Differential gene expression analysis based on the negative binomial distribution. Use with subprocess.run(['Rscript', '-e', 'library(DESeq2); ...']). |

**Table 29:** Continue software descriptions

| Software Name | Description |
|---|---|
| clusterProfiler | [R Package] A package for statistical analysis and visualization of functional profiles for genes and gene clusters. Use with subprocess calls. |
| DADA2 | [R Package] A package for modeling and correcting Illumina-sequenced amplicon errors. Use with subprocess calls. |
| xcms | [R Package] A package for processing and visualization of LC-MS and GC-MS data. Use with subprocess calls. |
| FlowCore | [R Package] Basic infrastructure for flow cytometry data. Use with subprocess calls. |
| edgeR | [R Package] Empirical Analysis of Digital Gene Expression Data in R, for differential expression analysis. Use with subprocess calls. |
| limma | [R Package] Linear Models for Microarray Data, for differential expression analysis. Use with subprocess calls. |
| harmony | [R Package] A method for integrating and analyzing single-cell data across datasets. Use with subprocess calls. |
| WGCNA | [R Package] Weighted Correlation Network Analysis for studying biological networks. Use with subprocess calls. |
| samtools | [CLI Tool] A suite of programs for interacting with high-throughput sequencing data. Use with subprocess.run(['samtools', ...]). |
| bowtie2 | [CLI Tool] An ultrafast and memory-efficient tool for aligning sequencing reads to long reference sequences. Use with subprocess.run(['bowtie2', ...]). |
| bwa | [CLI Tool] Burrows-Wheeler Aligner for mapping low-divergent sequences against a large reference genome. Use with subprocess.run(['bwa', ...]). |

**Table 30:** Continue software descriptions

| Software Name | Description |
|---|---|
| bedtools | [CLI Tool] A powerful toolset for genome arithmetic, allowing operations like intersect, merge, count, and complement on genomic features. Use with subprocess.run(['bedtools', ...]). |
| macs2 | [CLI Tool] Model-based Analysis of ChIP-Seq data, a tool for identifying transcript factor binding sites. |
| fastqc | [CLI Tool] A quality control tool for high throughput sequence data. Use with subprocess.run(['fastqc', ...]). |
| trimmomatic | [CLI Tool] A flexible read trimming tool for Illumina NGS data. Use with subprocess.run(['trimmomatic', ...]). |
| mafft | [CLI Tool] A multiple sequence alignment program for unix-like operating systems. Use with subprocess.run(['mafft', ...]). |
| Homer | [CLI Tool] Motif discovery and next-gen sequencing analysis. |
| FastTree | [CLI Tool] Phylogenetic trees from sequence alignments. |
| muscle | [CLI Tool] Multiple sequence alignment tool. |
| plink2 | [CLI Tool] A comprehensive toolkit for genome association studies that can perform a range of large-scale analyses in a computationally efficient manner. Use with subprocess.run(['plink2', ...]). |
| gcta64 | [CLI Tool] Genome-wide Complex Trait Analysis (GCTA) tool for estimating the proportion of phenotypic variance explained by genome-wide SNPs and analyzing genetic relationships. Use with subprocess.run(['gcta64', ...]). |
| iqtree2 | [CLI Tool] An efficient phylogenetic software for maximum likelihood analysis with built-in model selection and ultrafast bootstrap. Use with subprocess.run(['iqtree2', ...]). |

**Table 31:** Completeness scoring rubric for experimental documentation.

| Score | Level | Description |
|---|---|---|
| 1 | Severely Incomplete | Missing many major components necessary for experiments. Cannot be followed for implementation. Lacks critical methodological details and equipment specifications. |
| 2 | Significantly Incomplete | Missing few major components. Requires substantial additional effort from user. Key experimental parameters or control measures absent. |
| 3 | Moderately Complete | Contains basic components with some detail gaps. Requires moderate additional effort from user. Most critical parameters defined but needs optimization. |
| 4 | Mostly Complete | Contains nearly all necessary components. Requires minimal additional effort from user. Well-described procedures with adequate troubleshooting guidance. |
| 5 | Completely Thorough | Contains all necessary components in appropriate detail. Can be followed precisely with no additional effort. Comprehensive methodology, specifications, and contingency plans. |

**Table 32:** Accuracy scoring rubric for experimental documentation.

| Score | Level | Description |
|---|---|---|
| 1 | Severely Inaccurate | Contains many major errors or misconceptions. Methods would lead to invalid or unreliable results. Fundamental scientific principles violated or misapplied. |
| 2 | Significantly Inaccurate | Contains few major errors or several minor errors. Some methodological approaches are flawed. May lead to partially valid but questionable results. |
| 3 | Moderately Accurate | Generally correct but contains some minor errors or imprecisions. Core methodology is sound but may have optimization issues. Results would be acceptable but not optimal. |
| 4 | Mostly Accurate | Contains minimal errors. Methods align with accepted scientific practices. Results would be reliable with only minor corrections needed. |
| 5 | Completely Accurate | Contains no errors. Methods perfectly align with best scientific practices. Results would be highly reliable and reproducible. Includes appropriate controls and validation steps. |
